# A Novel mTOR-Stat3-Stathmin Pathway Establishes Oocyte Polarization Competence by Orchestrating Centrosome Regulation and Microtubule Organization

**DOI:** 10.1101/2025.02.27.640508

**Authors:** Amal Shawahny, Yoel Bogoch, Yaniv M. Elkouby

## Abstract

In animals and plants, egg production is essential for fertility, reproduction, and embryonic development. The production of functional eggs in sufficient numbers for a female lifespan requires dynamic and precisely coordinated cellular and developmental programs in early oogenesis. However, our understanding of their underlying regulatory mechanisms is critically lacking. Here, we aimed to identify overlooked essential regulators of early oogenesis in zebrafish. First, we present the establishment of a long-term ovary culture system which enables physiological oocyte development from oogonia to primordial follicles in cultured ovaries, providing an invaluable *ex-vivo* platform for rapid investigation. Next, we utilized this system for robust functional screening of candidates from stage-specific oocyte transcriptomic data. We identified mTOR, Stat3, and Stathmin as novel regulators of oocyte polarity. By using a combination of genetics, pharmacological manipulations *ex-vivo*, and rescue experiments, we established an essential mTOR-Stat3-Stathmin pathway that orchestrates centrosome regulation and microtubule organization to facilitate oocyte polarity. Microtubules control the localization and condensation of the essential polarity regulator Bucky ball (Buc). Loss of mTOR or Stat3 functions, as well as overactivation of Stathmin (a microtubule destabilizing protein) resulted in aberrant centrosome regulation and destabilization of microtubules, leading to dispersed mis-localized Buc condensates and loss of polarity. We show that mTOR acts upstream of Stat3 in oocytes, and that inhibition of Stathmin in *stat3^-/-^*or mTOR deficient ovaries rescued both cytoskeletal and polarity defects. We propose a novel mTOR-Stat3-Stathmin pathway which through cytoskeletal regulation, provides oocytes with polarization competence, a step likely widely conserved in biology. mTOR, Stat3, and Stathmin are known for their roles in cellular growth and cancer. Our work reveals their novel unpredicted functions in cell polarity during animal post-embryonic development.

## Introduction

In sexually reproducing species, egg production is essential for fertility, reproduction, and embryonic development. Early oogenesis requires dynamic and precisely coordinated cellular and developmental programs that are critical for egg production and reproduction. In mammals, early oogenesis occurs during fetal development and determines the number and quality of eggs for the entire lifespan. In humans, deficiencies in these early processes are the leading cause for infertility, miscarriages, and gonadal tumors,^1^ yet the underlying mechanistic defects remain unclear due to the lack of a complete understanding of these natural processes.

While obvious ethical and technical constrains limit investigation in humans, zebrafish developing ovaries provide an excellent model for early oogenesis.^2–5^ Ovaries in zebrafish are highly accessible experimentally, and oogenesis is conserved in a manner similar to that in mammals.^2^ To bridge the gap in our knowledge of early oogenesis, we aimed to identify novel regulators of early oocyte development, as stepping stones for uncovering essential mechanisms. We developed a novel long-term ovarian culture system as powerful *ex-vivo* platform for investigation, and used it to functionally screen for candidates from our stage-specific oocyte transcriptomic database. ^6^ This led us to uncover an unpredicted pathway which controls oocyte polarity.

Oocyte polarity is widely essential for embryonic development. In vertebrates, including zebrafish and *Xenopus*, the oocyte is polarized along an animal-vegetal axis, with the vegetal pole harboring key embryonic regulators.^77^ Embryonic dorsal- and germline fate determinants are organized as RNA-protein (RNP) complexes in specialized germ granules during oogenesis.^7^ Those determinants must localize to the oocyte vegetal pole, from which they later establish the global embryonic body axes and the germline lineage.^7^ The formation, integrity, and polarized localization of these determinant containing germ granules require the Bucky ball (Buc) protein in fish^89,109^ and its homolog XVelo in frogs. ^11–12^

In zebrafish, during symmetry breaking at the onset of oocyte differentiation, Buc becomes polarized by localizing around the centrosome.^9^ Buc subsequently organizes granules in a nuclear cleft,^9^ where they assemble a membraneless organelle called the Balbiani body (Bb).^13^ Buc is an intrinsically disordered protein which undergoes phase-separation to drive the formation of the Bb through molecular condensation.^14,10^ Buc forms dynamic liquid-like condensates that gradually generate the mature solid-like Bb compartment.^10,14^ The mature Bb then translocates to the cytoplasmic membrane. At the cortex, the Bb docks, disassembles, and unloads its passenger RNP complexes, specifying this region of the cortex as the oocyte’s vegetal pole.^15,16^ Loss of Buc functions in *buc^-/-^* oocytes results in radially symmetrical eggs and early embryonic lethality. ^17,18^

Buc dynamics, Bb formation and oocyte polarity are orchestrated by microtubules.^910^ Buc polarization around the centrosome at symmetry breaking requires microtubules, which are organized from the centrosome microtubule organizing center (MTOC).^9,10,19,4^ This microtubule organization simultaneously controls essential chromosome pairing dynamics during meiosis, forming the chromosomal bouquet - a configuration conserved from yeast to mammals.^9,4^ The bouquet centrosome MTOC machinery couples these key events, serving as a cellular organizer that forms an essential cilium for oocyte development, controls meiotic chromosomal pairing, and breaks the oocyte symmetry.^9^ ^19^ The subsequent localization of small Buc granules into the main condensate in the cleft requires dynein-mediated transport along microtubules.^10^ Later, microtubules are also essential for maintaining the integrity of the main condensate and for “caging” the mature Bb to maintain its size and shape.^10^

These mechanisms are widely conserved. The Bb is conserved in oocytes from insects to humans, exhibiting similar morphology and organelle content, and cellular and developmental dynamics.^13,20,21–24^ While the functions of the Bb in human oogenesis are still unclear, in mice, the Bb is associated with the proper formation of the primordial follicle,^22^ suggesting a critical role. In mice oocytes, Bb components localize around the centrosome and require microtubule trafficking.^22,25^ In the insect *Thermobia*, Bb components polarize and align with the bouquet configuration,^26^ likely via the conserved bouquet MTOC machinery. In *Drosophila* oocyte polarity RNP complexes in pole granules are organized by the Oskar protein through molecular condensation.^27,28^ Oskar and Buc are proposed to play analogous roles in granule organization,^11^ and Oskar exhibits similar condensation dynamics and trafficking over microtubules.^27,29,30^ However, despite the essential roles of microtubules in these processes, how their organization is regulated in early vertebrate oocytes to facilitate polarization remains unknown.

Here, we identified mTOR, Stat3, and Stathmin as novel regulators of zebrafish oocyte polarity. By using a combination of genetics, manipulations *ex-vivo*, and rescue experiments, we established an unpredicted mTOR-Stat3-Stathmin pathway that controls centrosome MTOC regulation and microtubule organization to facilitate oocyte polarity. Loss of mTOR or Stat3 functions, as well as overactivation of Stathmin (a microtubule destabilizing protein) resulted in abberrant centrosome regulation and destabilization of microtubules, leading to dispersed mis-localized Buc granule condensates and loss of polarity. By deciphering their hierarchical activities, we propose a novel mTOR-Stat3-Stathmin pathway that provides oocytes with polarization competence. Given that cellular polarization generally requires precise organization of cytoskeletal machineries, our findings are likely widely relevant in biology. Furthermore, while mTOR and Stat3 are known for their roles in cellular growth, proliferation, and metabolism, and Stat3 and Stathmin are involved in regulating abnormal centrosomes in cancer, our work reveals their novel functions in cell polarity during animal development in physiological *in-vivo* settings.

## Results

### A novel *ex-vivo* system to study early oogenesis and ovarian development in zebrafish and as a platform to identify novel regulators

Early oogenesis in zebrafish occurs during post-embryonic development in juvenile fish. Investigations of zebrafish ovaries thus far have relied on demanding genetic approaches. Unbiased identification of novel regulators has been possible through extensive four-generation maternal-effect forward genetic screens (e.g.,^18,31,32^). In such screens, recovering alleles that directly affect early oogenesis is challenging, as essential factors are detrimental to oogenesis. Furthermore, in zebrafish, severe defects in oogenesis often result in sex conversion, leading to the development of sterile males,^33–36^ which either challenges or completely precludes investigation of ovaries. Alternative reverse genetic approaches are also demanding, as factors that play crucial roles in oogenesis may also be involved in early embryonic development and/or other tissues. Consequently, the generation of mutants may lead to embryonic or larval lethality and require sophisticated conditional approaches for ovarian analyses. To circumvent these issues, we aimed to develop an *ex-vivo* system for the study of early oogenesis, providing a robust assay for testing the functions of new candidates before embarking on genetic studies.

Culture of zebrafish juvenile ovaries is possible for several hours in HL-15-based media and low-melt agarose embedding, and has been successful for live imaging experiments over the course of several hours.^10,19,37,38^ We expanded this system to develop a mounting-media-free system, along with conditions for culturing ovaries for several days (Fig. S1A). We optimized the mounting-media-free culture, which maintains ovary shape while allowing improved passage of material from the media. In the fish body, ovaries are held straight by connective tissues, and upon dissection, free live ovaries tend to curl up. To preserve the ovaries in their physiological morphology, we placed them in poly-D-Lysine (PDL)-treated glass-bottom dishes and covered them with a cell strainer mesh to hold them in a straightened and flattened position, interfacing with the glass bottom of the dish (Fig. S1A). We then optimized the culture media by replacing HL-15 with DMEM, which is richer in nutrients, and by adding supplements, including ovary extract, FBS, and fish serum, as sources of physiologically derived growth factors (basic ovary culture media, BOCM; Methods), and cultured the ovaries at 28°C for 10 days.

Ovaries cultured in BOCM remained viable for 7 days in culture, as shown by the vital mitochondrial dye Mitotracker, which selectively labels active mitochondria as a proxy for cell viability. Ovaries remained viable through 7 days post-culture (dpc), but were necrotic at 10 dpc (Fig. 1A-B). To evaluate oocyte development in culture, we pulsed ovaries with EdU at 1 dpc and chased for labeled cells at 7 dpc. EdU labeling was detected in the nuclei of cells in pulsed ovaries but not in EdU non-treated ovaries (Fig. 1C), confirming signal specificity. Co-labeling with the germ cell-specific marker Ddx4 showed that both Ddx4-positive germ cells and Ddx4-negative somatic cells were labeled with EdU, demonstrating that both somatic and germ cells divide in culture (Fig. 1D). However, all EdU-labeled Ddx4-positive germ cells were at the oogonia stage, and none were detected at meiotic stages (Fig. 1D, H). Therefore, while these conditions allow for ovary viability and cell division, they are insufficient to initiate and progress through oocyte differentiation and meiosis.

**Figure 1.**
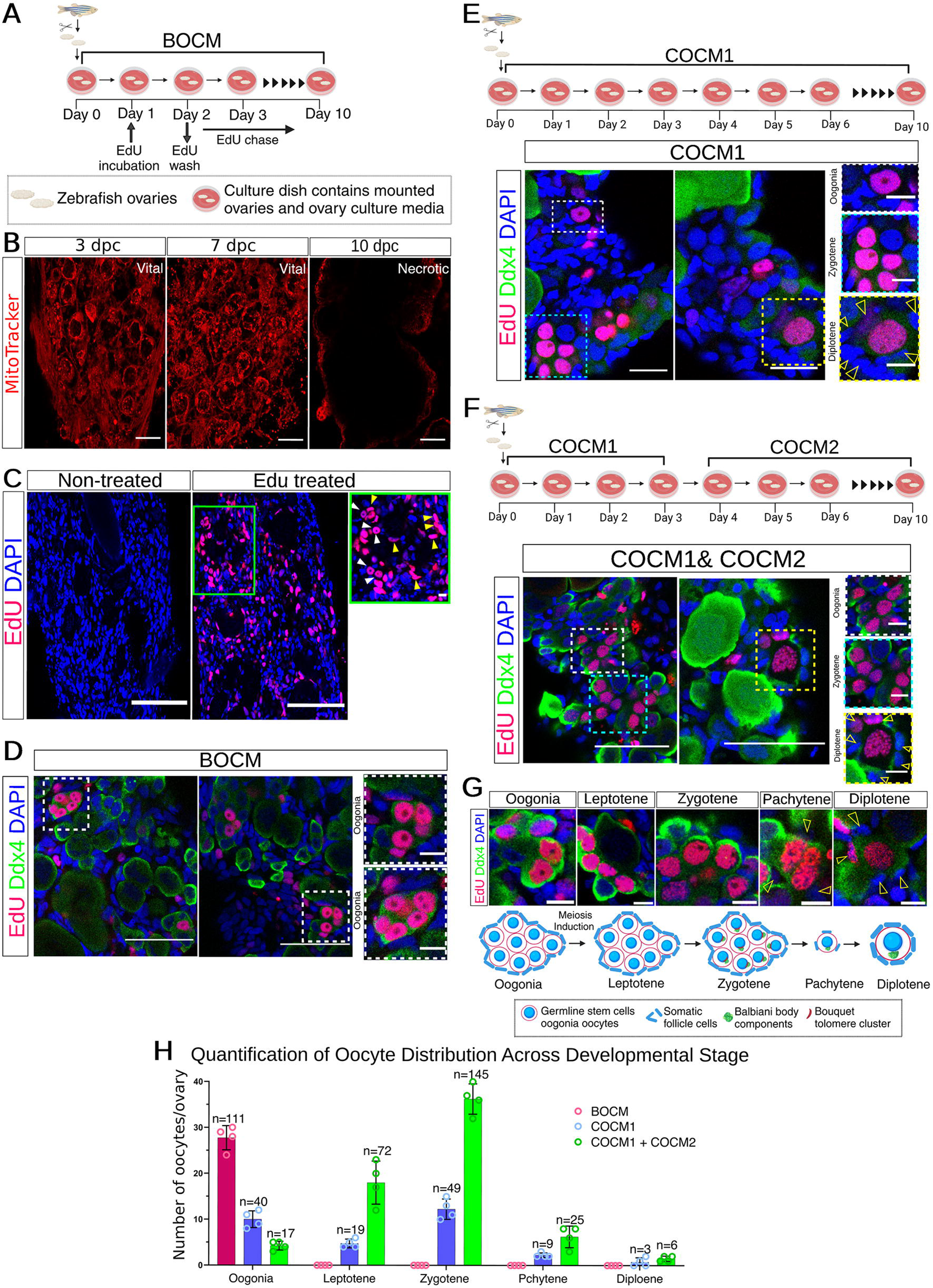
Long-term ovary culture as a novel *ex-vivo* system for investigation of early oogenesis and ovarian development. **A.** Experimental set up of ovary culture for 10 days in BOCM. EdU pulse and chase timepoints are indicated. **B.** Ovaries are vital for 7 but not 10 days in culture in BOCM. Cultured ovaries were live-labeled for Mitortracker at the indicated time points. Clear vital Mitotracker signals are detected through 7 dpc, but not in 10 dpc. n=4-7 ovaries per time-point. Scale bars are 40 μm. **C.** Cultured ovaries execute somatic- and germ cell divisions. EdU-positive germ cells (white arrowheads) and somatic cells (yellow arrowheads) are detected at 3 dpc in ovaries treated with EdU between 0-1 dpc (two right panels), but not in EdU non-treated ovaries (left) as control. Right panel is a zoom-in image of the green box in the middle panel. Ovaries are co-labeled for DAPI (blue). n=4 ovaries. **D.** Culture in BOCM supports cell division but not progression in oogenesis beyond oogonia. Cultured ovaries as in A-C were co-labeled for Ddx4 (green) EdU (red), and DAPI (blue). Both Ddx4+ germ cells and Ddx4(-) somatic cells divide in culture, and germ cells are detected at oogonia stages, but not in meiotic stages. Two representative overviews are shown. Right panels are zoom-in images of the green box in the middle panel, showing oogonia in germline cysts. n=6 ovaries. **E.** Culture in COCM1 media supports meiotic entry and progression. Top: Experimental set up of ovary culture on COCM1. Bottom: Cultured ovaries in COCM1 were co-labeled for Ddx4 (green) EdU (red), and DAPI (blue). EdU and Ddx4 double positive cells were detected through early diplotene stages. Two representative overviews are shown. Right panels are zoom-in images of color coded boxes in the left images, showing cysts of oogonia and zygotene oocytes, and a primordial follicle at diplotene as indicated. Arrowheads indicate nuclei of granulosa cells. n=6 ovaries. **F.** Culture in COCM1+COCM2 media supports efficient progression in meiosis and oogenesis from oogonia to primordial follicles. Top: Experimental set up of ovary culture on COCM1+COCM2. Bottom: Cultured ovaries in COCM1+COCM2 were co-labeled for Ddx4 (green) EdU (red), and DAPI (blue). EdU and Ddx4 double positive cells were detected through early diplotene stages. Two representative overviews are shown. Right panels are zoom-in images of color coded boxes in the left images, showing cysts of oogonia and zygotene oocytes, and a primordial follicle at diplotene as indicated. Arrowheads indicate nuclei of granulosa cells. n=6 ovaries. **G.** Representative Zoom-in images of oocytes from the ovaries in F, at all oocytes stages from oogonia through diplotene as indicated and as shown in the outline of early oogenesis below. A cyst of several cells is shown for oogonia, leptotene, and zygotene stages. Single oocytes in primordial follicles are shown for pachytene and diplotene stages Arrowheads indicate nuclei of granulosa cells. **H.** Frequencies of EdU labeled oocytes at the indicated stages under culture conditions in BOCM, or COCM1, or COCM1+COCM2 from experiments in D-G. Bars are mean±SD. In B-G, scale bars are 50 μm in zoom-out panels, and 10 μm in zoom-in panels.

We hypothesized that meiosis induction and oocyte differentiation in culture would require the activity of female hormones, and therefore added Estradiol-17 beta (E_2_) and 17alpha, 20beta-dihydroxy-4-pregnen-3-one (DHP) to our culture media (complete ovary culture media, COCM1; Methods). E_2_ induces oogonial proliferation^39^ and DHP acts directly on the initiation of the first meiotic division later in oogenesis.^39^ As shown by EdU pulse-chase experiments, EdU-labeled Ddx4-positive oocytes under these conditions were detected at oogonia, but also at leptotene and zygotene stages of meiosis I, with few continuing to pachytene and diplotene stages in primordial follicles (Fig. 1E, H). In addition, ovaries cultured in COCM1 remained vital through 10 dpc (Fig. S1B).

To improve our culture conditions for more efficient oocyte development, we optimized the time and duration of exposure to E_2_ and DHP throughout culture. Previous studies have shown that E_2_ can inhibit germline cyst breakdown,^40^ which might explain the low frequency of primordial follicles and oocytes at meiotic stages in ovaries cultured in COCM1. Therefore, attempting to mimic natural developmental and physiological regulation, we optimized our culture conditions such that both E_2_ and DHP (COCM1; Methods) are supplemented during 1-3 dpc to promote early development, while only DHP (COCM2; Methods) is supplemented during 4-10 dpc (Fig. 1F), to allow further differentiation through meiosis.

Indeed, under these conditions, we detected a significantly higher number of EdU-labeled Ddx4-positive oocytes at leptotene and zygotene stages, as well as an increased number of pachytene-stage oocytes in primordial follicles (Fig. 1F-H). However, the number of diplotene-stage oocytes was slightly increased but remained low (Fig. 1H). Further optimization will be needed in the future to enhance development from pachytene to diplotene stages. Nevertheless, our experiments demonstrate that our culture conditions (COCM1+COCM2) efficiently support early oogenesis from oogonia through pachytene stages and from the germline cyst to the formation of primordial follicles (Fig. 1G-H). Developing oocytes in culture showed both cyst and follicle formation dynamics, with normal morphology as well as co-organization of somatic cells, and appeared indistinguishable from oocytes in ovaries *in-vivo*. Thus, we conclude that our novel long-term ovarian culture reliably recapitulates early oogenesis and can be used as a powerful *ex-vivo* system for investigation.

### Transcriptomic-based screen in ovaries *ex-vivo* identifies Stat3 as a novel regulator of oocyte polarity in zebrafish oogenesis

To identify novel candidate regulators of zebrafish oogenesis, we mined our previously reported oogenesis transcriptomics database.^6^ This dataset includes the transcriptome of five major stages in early oogenesis, isolated by size: Symmetry breaking, Nuclear cleft, Mature Bb, St. II, and St. III.^2,6^ From ∼11,000 differentially expressed genes across these stages, we identified five distinct clusters of specific gene expression dynamics throughout oogenesis (Fig. 2A), as well as differentially expressed genes in pairwise comparisons between stages, providing a molecular characterization of oogenesis in zebrafish.^6^ Here, we mined this data in search of novel candidate regulators of zebrafish oogenesis. We formulated three criteria for candidate selection: ***1)*** Genes with unknown functions in zebrafish oogenesis to uncover new biological regulation, ***2)*** Genes with distinct timing of activity, indicating stage-specific developmental regulation, and ***3)*** Genes with available, established small molecule inhibitors, allowing for specific experimental manipulations.

**Figure 2.**
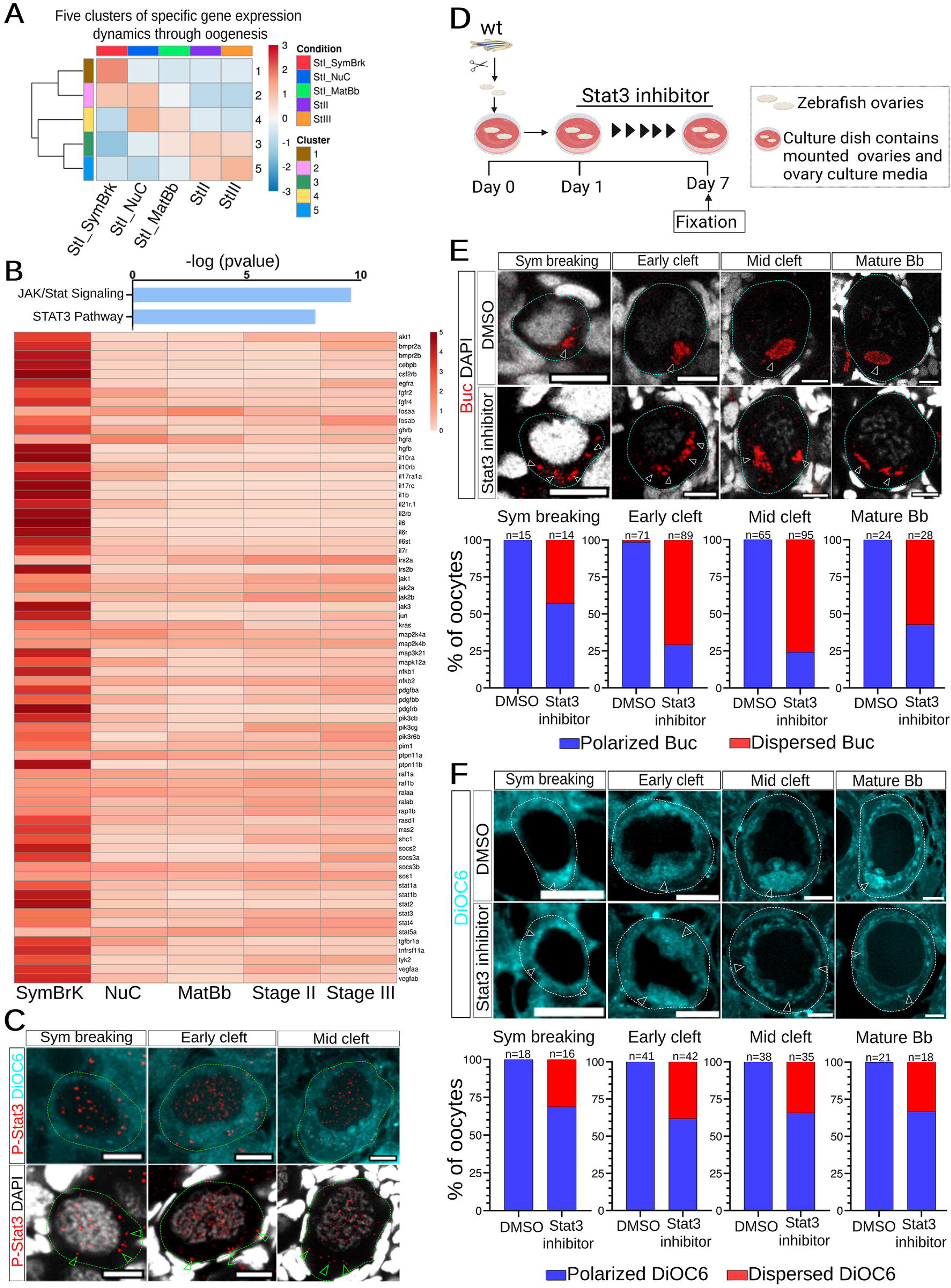
Identification of Stat3 as a novel regulator of oocyte polarity. **A.** Clusters of gene expression dynamics through five major stages in oogenesis from oocyte stage-specific transcriptomic data.^6^ Cluster #1 represent genes with enriched expression specifically in the St.I_SymBrk stage. **B.** Genes associated with the Stat3 and Jak/Stat pathways are expressed in oogenesis. Top: Stat3 and Jak/Stat pathways are enriched in cluster #1. Bottom: Expression levels through oogenesis of the genes included in Stat3 and Jak/Stat pathways from cluster #1. **C.** Stat3 is active in early oogenesis. p-Stat3 (red) labeling in ovaries co-labeled with DiOC6 (cyan, top), and DAPI (greyscale, bottom). p-Stat3 is detected in nuclei and cytoplasm (green arrowheads) of oocytes (dashed outlines) at symmetry breaking, early-, and late cleft stages. Same oocytes are shown with different channels in the top and bottom panels per stage. n=14 ovaries. **D.** Experimental set up of Stat3 inhibition in cultured ovaries *ex-vivo*. **E.** Inhibition of Stat3 *ex-vivo* perturbs Buc polarity. Representative images of oocytes (dashed outlines) at symmetry breaking, early-, and late cleft, and mature Bb stages in ovaries treated with DMSO (top) or Stat3 inhibitor (bottom) and labeled for Buc (red, white arrowheads) and DAPI (greyscale). The distribution of Buc phenotypes per stage is plotted below each panel. n=number of oocytes, from 6 DMSO treated and 6 Stat3 inhibitor treated ovaries. **F.** Inhibition of Stat3 *ex-vivo* perturbs the polarized localization of Bb-enriched organelles. Representative images of oocytes (dashed outlines) at symmetry breaking, early-, and late cleft, and mature Bb stages in ovaries treated with DMSO (top) or Stat3 inhibitor (bottom) and labeled for DiOC6 (cyan, white arrowheads). The distribution of DiOC6 phenotypes per stage is plotted below each panel. n= number of oocytes, from 6 DMSO treated and 6 Stat3 inhibitor treated ovaries. In all panels, scale bars are 10 μm.

We screened for candidate functions by applying their small molecule inhibitors in the novel *ex-vivo* ovary culture system. We isolated ovaries during early juvenile ovarian development from fish at 4 weeks post-fertilization (wpf). Wild-type (wt) ovaries were cultured for 7 days in the presence of the inhibitor of interest or vehicle as control. As a straightforward assay for general ovarian morphology, oocyte staging, and development, we labeled ovaries at the end of the culture with DAPI and the lipid dye DiOC6. DiOC6 labels membranes, including those of the nuclear envelope, organelles, and cytoplasmic membranes, and also shows some background in oocyte cytoplasm. Notably, DiOC6 also labels the enriched endoplasmic reticulum (ER) and mitochondria in the Bb, demarcating its stages of formation.^2,9,10^ Thus, DiOC6 is routinely used to visualize major cellular and morphological features of developing oocytes and ovaries.^2,9,10,19^ We initially scored DiOC6 phenotypes as part of our screening.

Among several enriched pathways that met our screening criteria, we identified the Stat3 pathway. In the canonical Stat3 pathway, several ligands and receptors can activate Stat3 through phosphorylation in a Jak-dependent or -independent manner (rev. in^41–44^). Active phosphorylated Stat3 (p-Stat3) translocates to the nucleus, where it transcriptionally regulates the expression of target genes.^41–44^ Cluster #1 in our transcriptomic data (Fig. 2A) includes genes whose expression is specifically higher in oocytes, corresponding to stages ranging from oogonia, leptotene, zygotene, and pachytene. This cluster includes genes active during early meiotic prophase, as well as symmetry breaking and early Buc nucleation in the nuclear cleft. Pathway analysis revealed an enrichment of transcripts for genes associated with both the Stat3 and JAK/Stat pathways in cluster #1 (Fig. 2B). This expression pattern suggests the activity of Stat3 and/or Jak/Stat at the appropriate developmental time window to control oocyte polarization. Supporting Stat3 activity during early oogenesis, we detected localization of p-Stat3 in the nuclei of oocytes from oogonia through diplotene (Fig. 2C). However, the role of Stat3 in zebrafish oogenesis had not been addressed previously.

We tested the function of Stat3 in oogenesis using its established small molecule inhibitor, 2,4-diamino-5-p-chlorophenyl-6-ethyl-pyrimidine (Pyrimethamine)^45^ in the *ex-vivo* ovary culture system. Towards screening ovaries *ex-vivo*, we first confirmed efficient inhibition by the relevant small molecule inhibitors. In case of Stat3, Pyrimethamine induces activity of Caspase proteins and apoptosis in human ovarian epithelial cancer cell line (A2780).^46^ We confirmed this in our hands and detected cleaved Caspase3 (cCaspase3) in A2780 cells treated with Pyrimethamine but not with control DMSO (Fig. S2). We next cultured 4 wpf wt ovaries for 7 days in the presence of Pyrimethamine or DMSO as a control (Fig. 2D). To ensure that ovaries from individual fish were at consistent developmental stages, we isolated ovaries from fish of a consistent standard length (SL; Fig. S3A). SL is a standardized measure for zebrafish post-embryonic development and also serves as an indicator of ovarian development.^2,47^

Ovaries in both Pyrimethamine- and DMSO-treated groups remained vital through culture, as indicated by normal Mitotracker signals (Fig. S3B), ruling out toxic effects of Pyrimethamine treatment. Both Pyrimethamine- and DMSO-treated ovaries also exhibited normal oocyte and ovarian morphology, with a typical distribution of oocyte stages, suggesting that Stat3 inhibition did not alter gross ovarian and oocyte development or induce major meiotic prophase progression defects (Fig. S3C). However, we observed specific polarity phenotypes upon Stat3 inhibition. Based on our studies of Bb formation in zebrafish, we established that Buc undergoes gradual condensation in forming the mature Bb condensate. We previously defined distinct steps in Bb formation^9^: symmetry breaking (oocytes at 10-15 µm in diameter), assembly in early nuclear cleft stages (oocytes at 17-25 µm in diameter), progression in mid-cleft stages (oocytes at 25-40 µm in diameter), and mature Bb (oocytes at 50-70 µm in diameter). We thus analyzed these defined steps in Bb formation in Stat3 deficient ovaries.

DMSO-treated control ovaries displayed normally polarized Buc granules in oocytes at symmetry breaking through cleft and mature Bb stages (Fig. 2E), as well as Bb-enriched mitochondria and ER, as labeled by the lipid dye DiOC6^9^ (Fig. 2F). In contrast, in Pyrimethamine-treated ovaries, Buc condensates were expanded and dispersed throughout the cytoplasm in oocytes at symmetry breaking (42.6%), early-(70.1%) and late (75.8%) cleft, as well as mature Bb (57.1%) stages (Fig. 2E). Similar phenotypes were detected for DiOC6 in 32.5-38.8% of oocytes throughout these stages (Fig. 2F). These results suggest polarity defects resulting from Stat3 inhibition.

To further rule out potential artifacts from the culture system, we confirmed these results in more physiological conditions by inhibiting Stat3 in whole, living fish. We treated 4 wpf fish with either Pyrimethamine or DMSO, adding the treatments to the fish water over two weeks of development (Fig. S4A). At the end of the treatment period, we sacrificed the fish and isolated their ovaries for analysis. To achieve efficient inhibition while minimizing potential stress and ensuring fish viability, we treated the fish with alternating cycles of treatment and rest, under standard husbandry conditions in our zebrafish facility (Methods). As a control for normal fish growth under these conditions, the SL of all fish in both groups was consistent before treatment at 4 wpf, and all fish reached similar and normal SL at 6 wpf (Fig. S4B), confirming that the experimental conditions did not affect general fish physiology and development.

Ovaries from DMSO-treated fish appeared normal, containing the typical distribution of oocyte stages and the expected single aggregates of Buc and DiOC6 (Fig. S4C-E). In contrast, 53.8% of ovaries from Pyrimethamine-treated fish were underdeveloped (Fig. S4C). These ovaries were smaller and contained fewer oocytes at the primordial follicle stage. However, oocytes that reached these stages in either underdeveloped or properly developed ovaries exhibited polarity defects, with ectopic and expanded Buc granules and DiOC6 signals (Fig. S4D-E), confirming the results from our culture experiments. The underdevelopment observed in Pyrimethamine-treated fish was specific and not seen in control-treated fish, nor was it observed in ovaries treated in culture, implying that this phenotype arose from indirect systemic effects of Stat3 inhibition in whole fish. Taken together, these *in-vivo* and *ex-vivo* results suggest that Stat3 is required for zebrafish oocyte polarity.

### *Stat3* mutant fish confirm that Stat3 is essential for oocyte polarity

To further test Stat3 function, we obtained fish carrying the *stat3* loss-of-function mutant allele *stat3^stl^*^27^ ^48^ and analyzed their ovaries. Loss of Stat3 in *stat3^stl^*^27^*^/stl^*^27^ fish (hereby referred to as *stat3^-/-^)* leads to lethality during juvenile post-embryonic development; however, mutant fish remain viable through 4 wpf^48^ (Fig. S5), which allowed us to examine ovarian development *in-vivo*. First, we detected the absence of nuclear phosphorylated Stat3 (p-Stat3) in oocytes in *stat3^-/-^* ovaries, compared to wt ovaries (Fig. 3A), confirming the lack of Stat3 activity in mutant ovaries, consistently with *stat3* loss-of-function phenotypes reported for this allele in embryonic development^48^ and regeneration.^49^ As reported, we found that *stat3* mutants develop scoliosis (Fig. S5A), and that their SL was slightly shorter that the SL of wt fish (Fig. S5B), likely because of their scoliosis phenotype^48^. However, their ovaries appeared to develop normally, containing a typical distribution of oocyte stages (Fig. 3B, Video S1), consistent with our *ex-vivo* studies.

**Figure 3.**
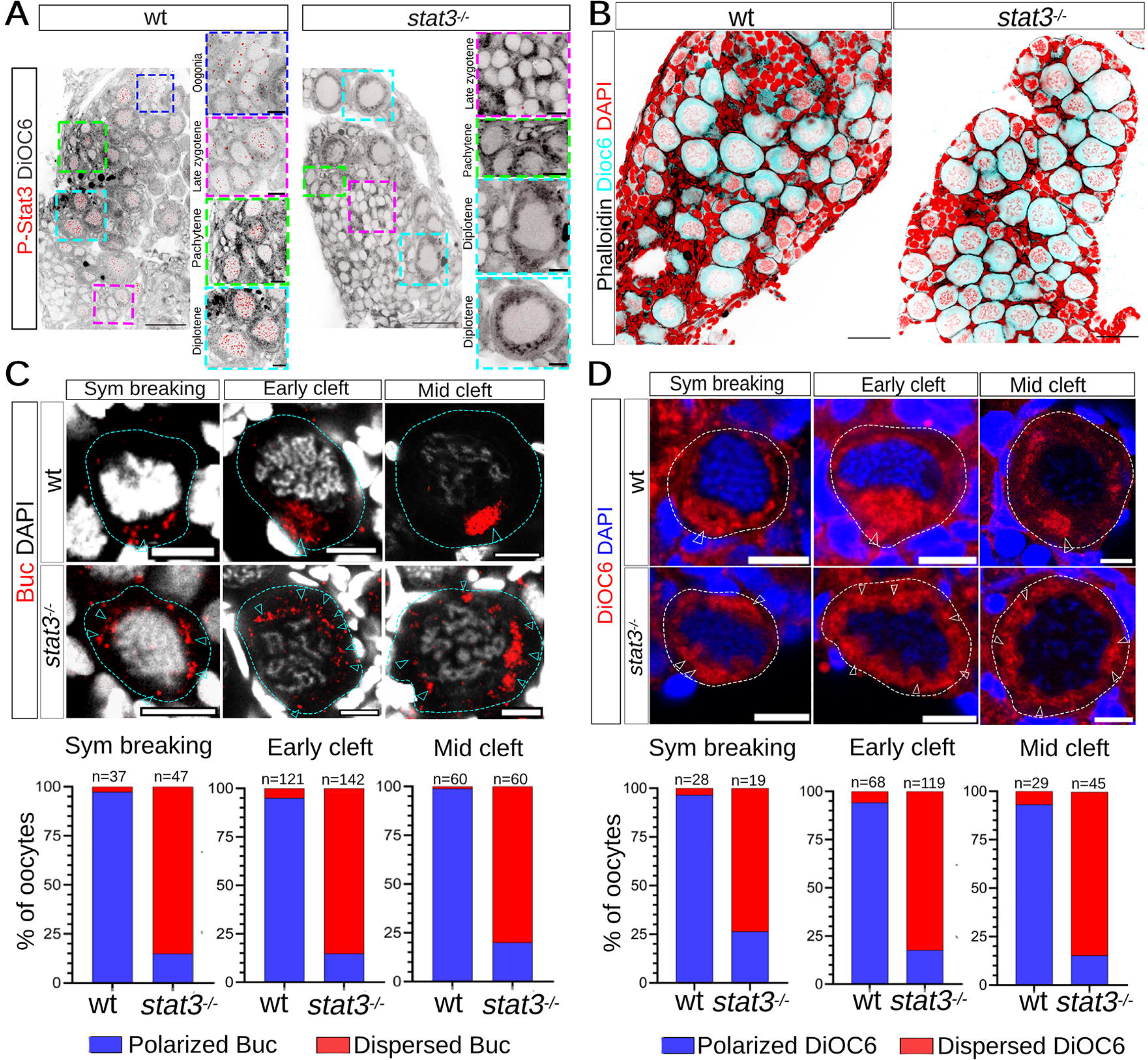
*stat3^-/-^* ovaries exhibit loss of oocyte polarity. **A.** p-Stat3 is not detected in *stat3^-/-^* ovaries, confirming stat3 loss of function. p-Stat3 (red) labeling in wt (left) and *stat3^-/-^*(right) ovaries, co-labeled with DiOC6 (inverted LUT). Right panels in each genotype are zoom-in images of the indicated oocytes stages from the color-coded boxes in the left panels. n=6 wt and 5 *stat3^-/-^* ovaries. Scale bars are 50 μm, and 10 μm in zoom-in panels. **B.** *stat3^-/-^*ovaries appear to develop normally and similar to wt. Representative images of wt (left) and *stat3^-/-^* (right) ovaries co-labeled for DiOC6 (cyan), Phalloidin (inverted LUT, black), and DAPI (red) are shown. n=6 wt and 6 *stat3^-/-^*ovaries. Scale bars are 50 μm. **C.** Stat3 is required for Buc polarization. Representative images of oocytes (dashed outlines) at symmetry breaking, early-, and late cleft stages in wt (top) and *stat3^-/-^* (bottom) ovaries labeled for Buc (red, cyan arrowheads) and DAPI (greyscale). The distribution of Buc phenotypes per stage is plotted below each panel. n= number of oocytes, from 8 wt and 7 *stat3^-/-^* ovaries. Scale bars are 10 μm. **D.** Stat3 is required for the polarized localization of Bb-enriched organelles. Representative images of oocytes (dashed outlines) at symmetry breaking, early-, and late cleft stages in wt (top) and *stat3^-/-^*(bottom) ovaries labeled for DiOC6 (red, white arrowheads) and DAPI (blue). The distribution of DiOC6 phenotypes per stage is plotted below each panel. n= number of oocytes, from 6 wt and 6 *stat3^-/-^* ovaries. Scale bars are 10 μm.

Strikingly, *stat3* mutant oocytes at all stages of Bb formation exhibited polarity defects (Fig. 3C-D). In wt ovaries, both DiOC6 and Buc showed normal polarized localization in symmetry-breaking through mid-cleft stages, as expected. In sharp contrast, in the majority of oocytes in *stat3* mutant ovaries the Bb appeared disassembled, with both Buc and DiOC6 signals dispersed and expanded in the cytoplasm, rather than being polarized (Fig. 3C-D). Oocytes at mature Bb stages (50-70 μm) are scarce in ovaries at 4 wpf, focusing our investigation on oocyte stages until mid-late cleft stages (40-50 μm). Nevertheless, we did observed few oocytes at mature Bb stages, and in *stat3^-/-^* ovaries they all showed a clear loss of Buc polarity, compared to wt (Fig. S6A), further confirming the polarity phenotypes in earlier stages.

Overall, based on our studies of Stat3 function using pharmacological tools both *ex-vivo* and in whole fish, as well as genetic analysis in *stat3* mutant fish, we conclude that Stat3 is essential for Buc localization, Bb formation, and oocyte polarity in zebrafish. We next addressed the mechanisms by which Stat3 controls Buc polarization.

### Stat3 is required for microtubule stabilization and regulation of the centrosome in oocytes

Stat3 is well known for its regulation of transcription,^41–44^ and could thus regulate the expression of an intermediate factor which in turn controls oocyte polarity. However, the specific polarity and Buc localization defects we observe upon loss of Stat3 functions are challenging to explain by direct transcriptional regulation alone. Interestingly, evidence for non-nuclear functions of Stat3 exist from cell culture models,^50–52^ as well as for its mitochondrial roles in cellular respiration in mice.^53^ In particular, Stat3 was shown to control microtubule stability independently of transcriptional regulation, by regulating the Stathmin protein.^50,51^ Stathmin disassembles microtubules by destabilizing the +end tips of microtubules, which results in their depolymerization, and by binding and sequestering free tubulin molecules, preventing their polymerization into microtubules.^54^ Stathmin can be found either in an active form cytoplasmically or can be deactivated by phosphorylation and shuttling to Golgi membranes.^54^ It has been established in cell culture models that Stat3 controls Stathmin activity by binding Stathmin’s tubulin interacting domain, and thereby antagonizing Stathmin microtubule destabilization activity.^50,54^ Thus, considering the essential roles that microtubules play in oocyte polarization and Bb formation,^9,10^ we investigated whether Stat3 controls microtubules through Stathmin regulation in the oocyte.

First, in addition to nuclear p-Stat3 signals, we detected puncta of p-Stat3 in oocyte cytoplasm (Fig. 2C, 3A). Cytoplasmic p-Stat3 signals were abolished together with the nuclear signals in *stat3^-/-^* ovaries (Fig. 3A), ruling out specificity issues and demonstrating that those puncta very likely represent phosphorylated Stat3 molecules. Cytoplasmic p-Stat3 puncta suggest that active Stat3 can be correctly poised for microtubule regulation in oocyte cytoplasm, consistently with previous reports of non-nuclear and cytoplasmic p-Stat3 functions.^50–52^

We tested whether loss of Stat3 function leads to microtubule instability. Examination of the general microtubule organization in ovarries via α-tubulin (αTub) labeling showed an elaborated network of microtubule cables and bundles throughout oocytes’ cytoplasm, including microtubules of the bouquet MTOC machinery at symmetry breaking,^9,10,19^ and those of the cleft meshwork in cleft stage oocytes^9,10,19^ (Fig. 4A). By contrast, in oocytes in *stat3^-/-^* ovaries, microtubules of the bouquet MTOC machinery and of the cleft meshwork were absent, and we detected much fewer cables in oocytes’ cytoplasm overall (Fig. 4A, Video S2), suggesting their depolymerization. Microtubules further appeared depolymerized in the few mature Bb stage oocytes detected at 4 wpf, in *stat3^-/-^*ovaries compared to wt (Fig. S6B). This observation was confirmed by comparing stabilized microtubule organization in wt and *stat3* ovaries, as labeled by acetylated-tubulin. Oocytes at various stages of Bb formation in wt ovaries exhibited a network of acetylated microtubules throughout the oocyte cytoplasm (Fig. 4B). However, acetylated microtubule signals were completely abolished in most oocytes in *stat3* ovaries (Fig. 4B). These results strongly suggest that Stat3 is essential for microtubule stabilization and organization in the oocyte.

**Figure 4.**
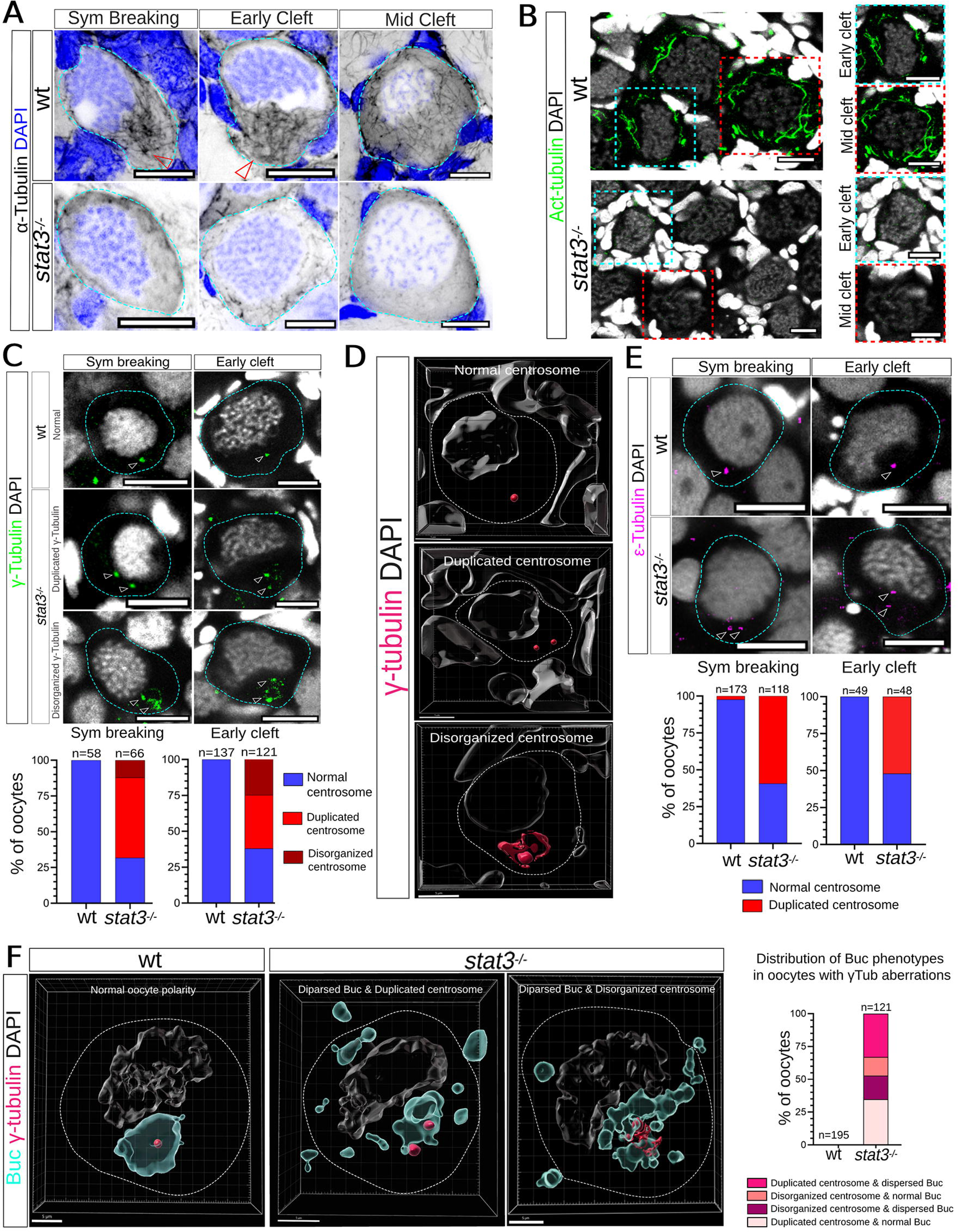
Stat3 is required for microtubule organization and centrosome regulation in oogenesis. **A.** Loss of microtubule organizations in *stat3* ovaries. Representative images of oocytes (dashed outlines) at symmetry breaking, early-, and late cleft stages in wt (top) and *stat3^-/-^* (bottom) ovaries labeled for αTub (Inverted LUT) and DAPI (blue). Red arrowheads indicate the bouquet MTOC machinery (top left panel) and the microtubule meshwork in the cleft (top mid panel) in the wt. n=10 wt and 7 *stat3^-/-^*ovaries. **B.** Microtubules are likely destabilized upon loss of stat3. Representative overview images of wt (top) and *stat3^-/-^* (bottom) ovaries labeled for acetylated tubulin (Act. Tub, green) and DAPI (greyscale). Right panels are zoom-in images of early- and mid-cleft stage oocytes from color-coded boxes in left panels. n=9 wt and 8 *stat3^-/-^* ovaries. **C.** Centrosome aberrations in *stat3* ovaries. Representative images of oocytes (dashed outlines) at symmetry breaking and early cleft stages in wt (top) and *stat3^-/-^* (middle and bottom panels) ovaries labeled for γTub (green) and DAPI (greyscale). In *stat3^-/-^* ovaries γTub (white arrowheads) phenotypes showed either two foci (mid panels) or cloud-like disorganized signals (bottom panels). The distribution of γTub phenotypes per stage is plotted below each panel. n=number of oocytes, from 9 wt and 7 *stat3^-/-^* ovaries. **D.** Representative 3D segmentation images of normal, duplicated, and disorganized γTub (maroon) phenotypes from E through entire oocyte volumes (dashed outlines). DAPI (greyscale) shows DNA of oocyte and surrounding cells. **E.** Centriole aberrations in *stat3* ovaries. Representative images of oocytes (dashed outlines) at symmetry breaking and early and cleft stages in wt (top) and *stat3^-/-^* (bottom) ovaries labeled for αTub (magenta) and DAPI (greyscale). In *stat3^-/-^* ovaries, αTub (white arrowheads) exhibited two foci. The distribution of αTub phenotypes per stage is plotted below each panel. n=number of oocytes, from 6 wt and 6 *stat3^-/-^*ovaries. **F**. Representative 3D segmentation images of oocytes from wt (left) and *stat3^-/-^* (middle and right panels) ovaries co-labeled for γTub (maroon), Buc (cyan), and DAPI (greyscale). 3D segmentation through entire oocyte volumes (dashed outlines) shows concomitant loss of Buc polarity with either γTub duplicated (mid panel) or cloud-like disorganized (right) signals. The distribution of Buc phenotypes in oocytes with γTub aberrations is plotted on the right. n=number of oocytes, from 5 wt and 4 *stat3^-/-^*ovaries. Scale bars are 10 μm in A-E and 5 μm in F.

In addition to microtubule stabilization, Stat3 is also known for its regulation of centrosomes in cancer, where it is essential for clustering abnormally duplicated centrosomes.^51^ Recently, Stat3 was proposed to control centrosome regulation in a transcriptional-independent manner, at least partly through Stathmin regulation.^51^ In zebrafish oogenesis, the centrosome microtubule-organizing center (MTOC) machinery is essential for both Buc polarization in symmetry breaking and Buc nucleation in early cleft stages,^9,10^ and the centrosome MTOC disassembles thereafter.^9^ We therefore investigated whether Stat3 is required for normal centrosome and MTOC regulation in zebrafish oogenesis.

The centrosome is composed of a pair of centrioles, which are surrounded by dozens of proteins that comprise the pericentriolar material (PCM).^55,56^ In centrosome maturation, PCM modifications lead to the recruitment of γTub, which in turn nucleates microtubules.^55,56^ We analyzed the localization of γTub in symmetry breaking and early cleft stage oocytes in *stat3^-/-^* ovaries. In wt oocytes, γTub localizes to the single centrosome at both symmetry breaking and early cleft stages (Fig. 4C). By contrast, γTub localization was abnormal in *stat3^-/-^* ovaries. At both symmetry-breaking stages and cleft stages, γTub was normally detected as a single centrosome in only 31.8% or 38% of oocytes, respectively (Fig. 4C). In most cases, γTub either formed two foci resembling two centrosomes, or was disorganized and appeared like a cloud (Fig. 4C). Both phenotypes were confirmed by three-dimensional (3D) segmentation through the entire oocyte volume (Fig. 4D, Video S3).

The observed γTub phenotypes could reflect defects in PCM and/or centriole regulation. To address whether centrioles are affected in *stat3^-/-^* ovaries, we labeled ovaries for the centriole marker αTub.^57^ As expected, oocytes in wt ovaries at both symmetry-breaking and cleft stages, exhibited a single focus of αTub signals, consistently with a single centrosome at these stages (Fig. 4E). However, most oocytes at both stages in *stat3^-/-^* ovaries exhibited two αTub foci (Fig. 4E), similar to γTub in *stat3* mutant ovaries. Thus, Stat3 is required for proper centriole and PCM regulation in zebrafish oogenesis. The ectopic foci of γTub and αTub may indicate roles for Stat3 in controlling centrosome duplication and/or centriole regulation.

Co-labeling of γTub and Buc in wt and *stat3^-/-^* ovaries confirmed the concomitant loss of polarity alongside abnormal centrosome regulation. 3D segmentation of entire oocyte volumes revealed that 48.8% of oocytes with either two foci of γTub or its cloud-like disorganization, also exhibit dispersed Buc condensates (Fig. 4F, Video S4). Altogether, these results strongly suggest that Stat3 is essential for microtubule stabilization and proper organization of the centrosome MTOC machinery in early oogenesis. They further propose that Stat3 may control oocyte polarity through the regulation of Stathmin.

### Stat3 acts upstream of Sathmin in controlling the centrosome MTOC machinery and oocyte polarity

Supporting our hypothesis that Stathmin is involved in oocyte polarity, we identified enrichment of a “Stathmin in breast cancer” pathway in our transcriptomic data, in the same cluster as the Stat3 and Jak/Stat pathways (Fig. 5A). While Stathmin’s functions are primarily studied in cancer biology, expression of genes associated with Stathmin in oocytes at the correct developmental timing, suggest it is involved in oocyte polarization. As further evidence, we detected p-Stathmin signals in oocytes (Fig. 5B). Detected p-Stathmin signals were prominent at symmetry breaking and early cleft stages and reduced at mid cleft stages (Fig. 5B), indicating dynamic Stathmin regulation in oocytes. Therefore, we tested whether Stat3 regulates oocyte polarity through Stathmin.

**Figure 5.**
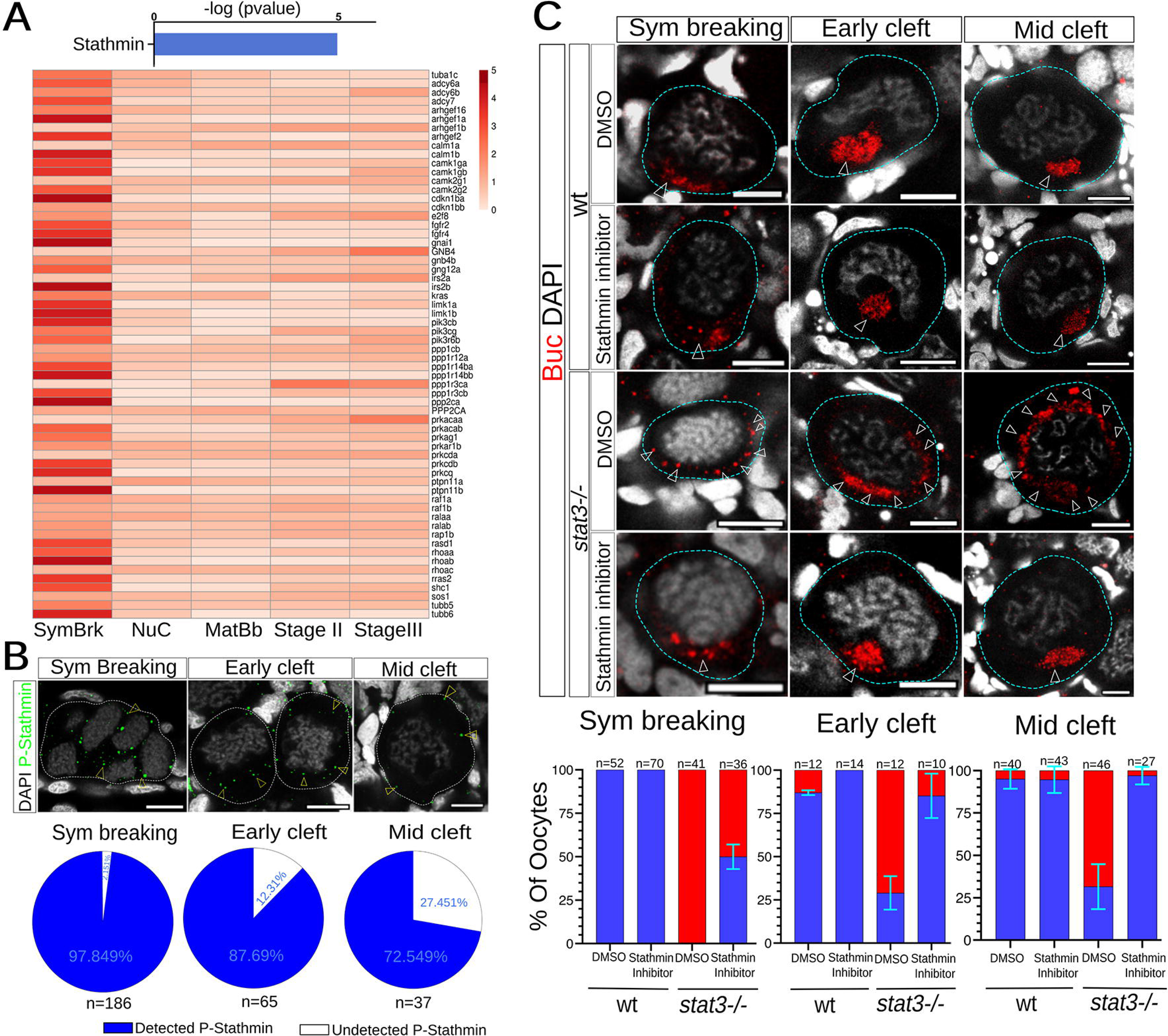
Stathmin functions in oogenesis and acts downstream of Stat3 in oocyte polarity. **A.** Genes associated with the Stathmin pathway are expressed in oogenesis. Top: the Stathmin pathway is enriched in cluster #1 (from Fig. 2A). Bottom: Expression levels through oogenesis of the genes included in the Stathmin pathway from cluster #1. **B.** p-Stathmin is detected in early oocytes. Representative images of oocytes at symmetry breaking (dashed line outlines a cyst of several oocytes), early-, and mid-cleft stages (dashed line outlines individual oocytes) from wt ovaries labeled for p-Stathmin (green, yellow arrowheads) and DAPI (greyscale). The frequencies of oocytes with detected p-Stathmin per stages are plotted below. n=number of oocytes, from 6 ovaries. Scale bars are 10 μm. **C.** Inhibition of Stathmin rescues Buc polarity in *stat3^-/-^* ovaries. wt (top two panels) or *stat3^-/-^*(bottom two panels) ovaries were cultured and each treated with DMSO as control, or Stathmin inhibitor, as indicated. Representative images of oocytes (dashed outlines) at symmetry breaking, early-, and mid-cleft stages from ovaries from each sample, labeled with Buc (red, white arrowheads) and DAPI (greyscale), are shown. Scale bars are 10 μm. The distribution of Buc phenotypes per stage is plotted below each panel. n=number of oocytes, from 4-5 ovaries sample. Error bars are mean±SD between experiments.

We hypothesize that normally in oocytes Stat3 antagonizes Stathmin, and upon Stat3 loss-of-function, Stathmin becomes overactive, destabilizing microtubules and leading to mis-localization of Buc condensates and loss of polarity. If this hypothesis is correct, inhibiting Stathmin in *stat3^-/-^*ovaries should prevent microtubule destabilization and rescue polarity. To test this, we used the small molecule Stathmin inhibitor GDP366,^58^ to inhibit Stathmin in *stat3^-/-^*ovaries. We cultured 4 wpf wt and *stat3^-/-^* ovaries, each treated with GDP366 or DMSO as control, and labeled them for Buc to monitor its localization (Fig. 5C). As expected, wt ovaries treated with GDP366 or DMSO exhibited normally polarized Buc condensates at symmetry breaking through cleft stages (Fig. 5C). Since microtubules are normal in wt ovaries, potential over-stabilization by Stathmin inhibition is unlikely to perturb polarity. Similar to uncultured *stat3^-/-^* ovaries above, cultured *stat3^-/-^* ovaries treated with DMSO showed mis-localized Buc in all or most oocytes throughout these stages (Fig. 5C). However, inhibition of Stathmin in *stat3^-/-^* ovaries treated with GDP366 restored Buc localization, with normally polarized condensates similar to wt (Fig. 5C). Levels of rescued polarity was observed from 50±7% of oocytes at symmetry breaking to 85±12.8% of oocytes at early cleft stages, and nearly complete rescue of 97±5.9% of oocytes at mid cleft stages (Fig. 5C).

The rescue of Buc polarity in *stat3* mutant ovaries upon Stathmin inhibition concludes that Stathmin acts downstream of and is likely regulated by Stat3. Stathmin inhibition likely prevents microtubule destabilization in the absence of Stat3 activity. To determine if this is the case, we examined microtubule organization in *stat3^-/-^* ovaries treated with Stathmin inhibitor in the same experimental setting above. We cultured 4 wpf wt and *stat3^-/-^*ovaries, each treated with GDP366 or DMSO as control, and labeled them for αTub.

As expected, we detected overall normal microtubule organization in wt ovaries treated with either GDP366 or DMSO, and lack of clear microtubule cables in *stat3^-/-^* ovaries treated with DMSO (Fig. 6A). However, in *stat3^-/-^* ovaries treated with GDP366, detection of clear microtubule cables and bundles was restored (Fig. 6A). Analyses of these phenotypes per stage during polarization at symmetry breaking through cleft stages confirmed rescue of microtubule organization in *stat3^-/-^* oocytes, by Stathmin inhibition (Fig. 6B). Normal microtubule organization was clear at all stages in wt ovaries treated with either GDP366 or DMSO, and their lack was prominent in most oocytes in *stat3^-/-^* ovaries treated with DMSO. By contrast, the majority of oocytes in *stat3^-/-^* ovaries treated with GDP366 exhibited the normal microtubule organization throughout these stages (Fig. 6B). Rescue levels ranged between 60% of oocytes at symmetry breaking, 79% at early-clef and 93.5% in mid-cleft oocytes (Fig. 6B).

**Figure 6.**
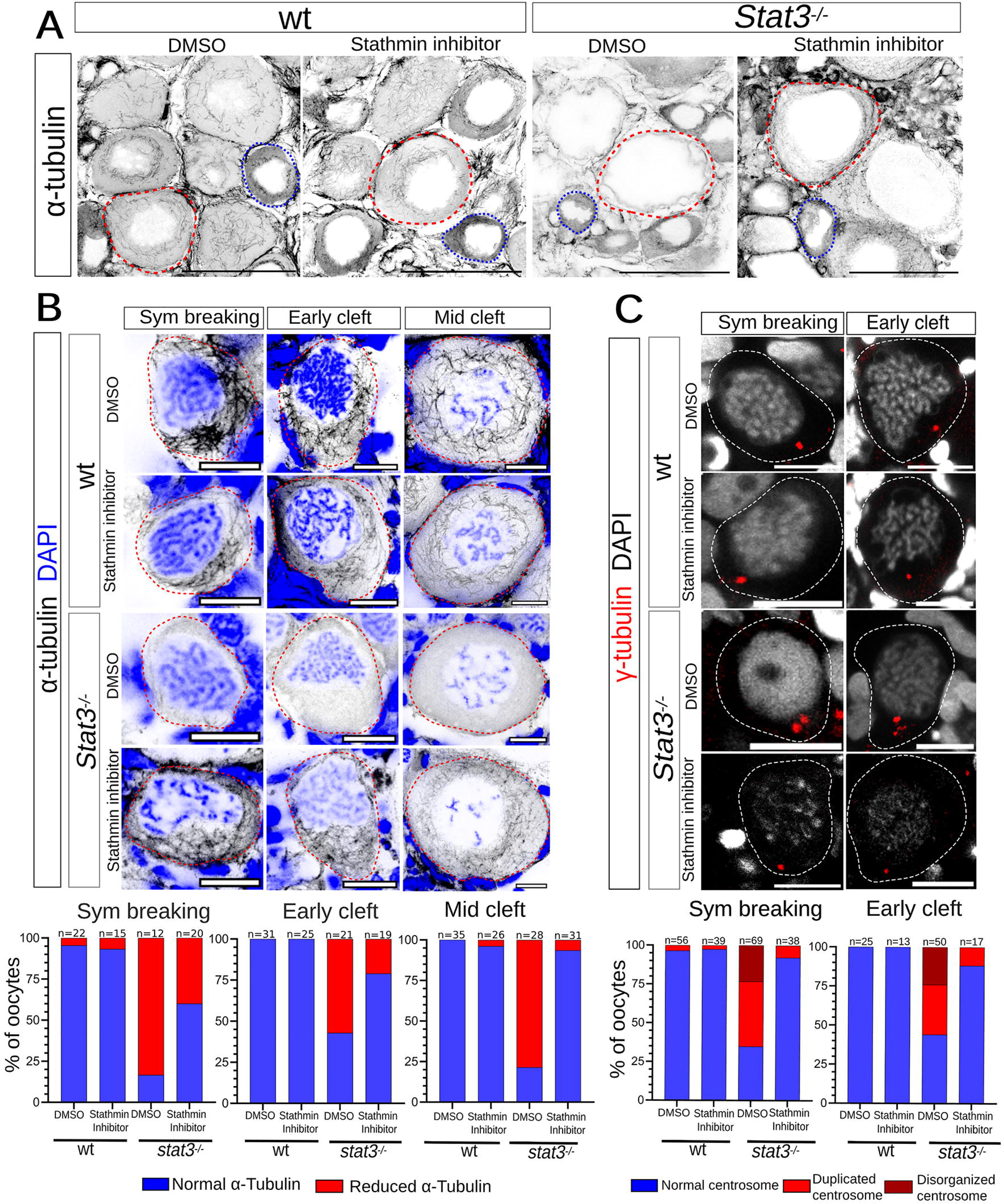
Stathmin regulates microtubule organization and the centrosome downstream of Stat3. A-B. Inhibition of Stathmin rescues microtubule organization in *stat3^-/-^*ovaries. wt or *stat3^-/-^* ovaries were cultured and each treated with DMSO, or Stathmin inhibitor, as indicated. Ovaries from each sample were labeled with αTub (inverted LUT) and DAPI (in B, blue). In A, overview images of wt (two left panels) and *stat3^-/-^*(two right panels) ovaries are shown (without DAPI channel). Outlines indicate oocytes at early- and mid-cleft (blue) and mature Bb (red) stages. In B, representative images of oocytes (dashed outlines) at symmetry breaking, early-, and mid-cleft stages from each wt (two top panels) and *stat3^-/-^* (two bottom panels) samples in A and treated with DMSO or Stathmin inhibitor as indicated are shown. The distribution of αTub phenotypes per stage is plotted below each panel. n= number of oocytes, from 4 ovaries per sample. **C.** Inhibition of Stathmin rescues γTub aberrations in *stat3^-/-^*ovaries. Representative images of oocytes (dashed outlines) at symmetry breaking and early cleft stages from ovaries labeled for γTub (red) and DAPI (greyscale) from similar experiments as in A-B. The distribution of γTub phenotypes per stage is plotted below each panel. n= number of oocytes, from 5-6 ovaries per sample. Scale bars are 50 μm in A, 10 μm in B-C.

We tested whether the centrosome phenotypes in *stat3^-/-^* ovaries can also be rescued by inhibition of Stathmin, by labeling for γTub in similar experiments. Normal single focus γTub-positive centrosomes were detected in oocytes at symmetry breaking and early cleft stages in wt ovaries treated with either GDP366 or DMSO (Fig. 6C). In *stat3^-/-^* ovaries treated with DMSO most oocytes at either stage exhibited either two foci or the cloud-like disorganized signals of γTub, as expected (Fig. 6C). Like microtubules, Stathmin inhibition in *stat3^-/-^* ovaries treated with GDP366 exhibited a normal single centrosome labeled with γTub, with rescue in 92.1% of oocytes at symmetry breaking and 88.2% of oocytes at early cleft stages (Fig. 6C).

Our results suggest that Stat3 is required to antagonize Stathmin, and that this regulation is crucial for microtubule organization, centrosome MTOC formation, and oocyte polarization. Stathmin likely fine-tunes the balance of microtubule stability and organization in oocytes. Given the importance of the centrosome MTOC machinery and microtubules in oocyte polarization, we propose that Stat3 plays a key role in maintaining a functional microtubule network, crucial for the localization of Buc. In this context, by ensuring correct microtubule organization, Stat3 provides the oocyte with the potential to polarize and thus functions as a polarity competence factor.

### Stat3 is regulated upstream by mTOR signaling

Having established the roles of Stat3-Stathmin in controlling microtubule organization in the oocyte, we next explored potential upstream regulators of Stat3. In our search for regulatory pathways, we identified two known activators of Stat3 - mTOR and IL-6^42–44,59^-which are enriched alongside Stat3 in our transcriptomic data (Fig. 7A, S7), and we focused on mTOR. mTOR pathway components have been demonstrated to play a role in sex determination in juvenile bipotential gonads in zebrafish,^60^ but its potential role in oocyte polarity had not been examined. The mTOR pathway is part of an intricate network of signaling pathways, making functional studies of mTOR specifically in oocyte polarity by genetics *in-vivo* challenging. To address this, we tested the role of mTOR by inhibiting its activity in ovaries *ex-vivo*.

**Figure 7.**
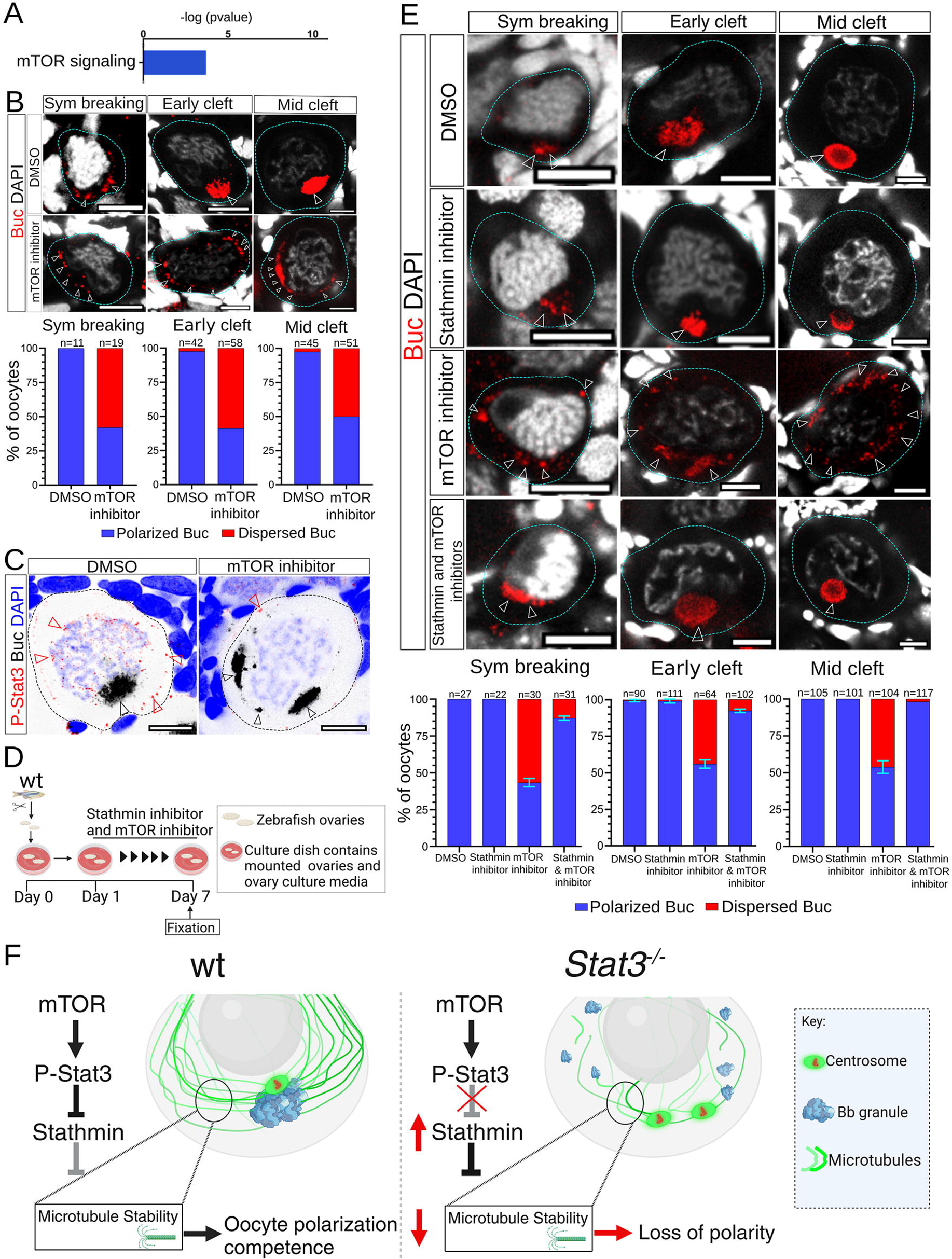
mTOR is required for Buc polarity and acts upstream of Stat3-Stathmin. **A.** Genes associated with the mTOR pathway are expressed in oogenesis. The mTOR pathway is enriched in cluster #1 (from Fig. 2A). **B.** mTOR is required for Buc polarity. Representative images of oocytes (dashed outlines) at symmetry breaking, early-, and mid-cleft stages in ovaries treated with DMSO (top) or mTOR inhibitor (bottom) and labeled for Buc (red, white arrowheads) and DAPI (greyscale). The distribution of Buc phenotypes per stage is plotted below each panel. n=number of oocytes, from 6 DMSO-treated and 6 mTOR inhibitor treated ovaries. **C.** mTOR is required for activation of Stat3. Representative images of oocytes (dashed outlines) in ovaries treated with DMSO (left) or mTOR inhibitor (right) and co-labeled for p-Stat3 (red, red arrowheads), Buc (inverted LUT, black arrowheads) and DAPI (blue). Inhibition of mTOR results loss of p-Stat3 detection in ovaries. n=4 DMSO treated and 4 mTOR inhibitor treated ovaries. Loss of p-Stat3 was detected in the same oocytes with loss of Buc polarity as shown. **D.** Experimental set up of rescue experiments in E: inhibition of mTOR and/or Stathmin in cultured ovaries *ex-vivo*. **E.** Inhibition of Stathmin rescues Buc polarity in mTOR deficient ovaries. wt ovaries were cultured and each treated with DMSO as control, or Stathmin inhibitor, or mTOR inhibitor, or both mTOR and Stathmin inhibitors, as indicated. Representative images of oocytes (dashed outlines) at symmetry breaking, early-, and mid-cleft stages in ovaries from each sample, labeled with Buc (red, white arrowheads) and DAPI (greyscale), are shown. The distribution of Buc phenotypes per stage is plotted below each panel. n=number of oocytes, from 6 ovaries per sample. Error bars are mean±SD between experiments. Scale bars in B, C, E are 10 μm. **F.** A novel mTOR-Stat3-Sathmin pathway establishes oocyte polarization competence in early oogenesis. Proper organization of microtubules and the centrosome MTOC is essential for oocyte polarity. In normal wt conditions (left), mTOR activates Stat3, which antagonizes and inhibits Stathmin. Stathimn destabilizes microtubules, and its inhibition by Stat3 ensures an optimally stabilized organization of microtubules and the centrosome MTOC, for potentiating oocyte polarization. mTOR may activate State3 in cooperation with other signaling molecules (e.g., Il-6). In *stat3^-/-^* ovaries (right), loss of Stat3 function leads to Stathmin overactivity, which destabilizes microtubules and perturbs centrosome regulation. Under these conditions, without proper microtubule and centrosome MTOC organizations, oocytes fail to polarize.

We cultured wt ovaries and treated them with the established small molecule inhibitor of mTOR, Rapamycin, or DMSO as a control, and assayed for Buc localization (Fig. 7B). In contrast to control-treated ovaries, inhibition of mTOR activity resulted in a loss of polarity, with Buc condensates mis-localized and dispersed throughout the cytoplasm in the majority of oocytes at symmetry breaking through cleft stages (Fig. 7B). These results suggest that mTOR is required for proper Buc polarity, phenocopying the effects observed in *stat3* mutant ovaries. Consistent with Stat3 activation by mTOR, we observed a reduction in nuclear and cytoplasmic p-Stat3 signals upon mTOR inhibition in ovaries, concomitantly with Buc polarity loss (Fig. 7C), indicating that mTOR is required for Stat3 activation in oocytes.

These results suggest that mTOR functions upstream of Stat3-Stathmin in regulating oocyte polarity. Based on this, we hypothesized that inhibition of Stathmin activity should rescue Buc polarity upon loss of mTOR function. To test this, we cultured wt ovaries *ex-vivo* and treated them with either the Stathmin inhibitor GDP366, Rapamycin, or a combination of both, side by side with DMSO as a control (Fig. 7D-E). As expected, treatment with DMSO or GDP366 alone resulted in normal Buc polarity at symmetry breaking and cleft stages (Fig. 7E). Treatment with Rapamycin alone led to a loss of Buc polarity, with mis-localized Buc condensates in oocytes throughout symmetry-breaking (56.6±2.8%), early cleft (44±2.8%), and mid-cleft (46.1±4.2%) stages (Fig. 7E). However, when GDP366 was added to Rapamycin-treated ovaries, a near-complete rescue of Buc polarity was observed in throughout these stages (87.1±1.4% at symmetry breaking, 92.3±1.1% at early cleft-, and 98.2±0% at mid-cleft stages; Fig. 7E). These results demonstrate that Stathmin activity is likely downstream of mTOR signaling.

Altogether, our work establishes a mTOR-Stat3-Stathmin pathway that regulates oocyte polarity during early oogenesis, revealing unanticipated functions in this process. In this pathway, mTOR signaling activates Stat3, and Stat3 is required for the inhibition of Stathmin (Fig. 7F). Ultimately, this pathway ensures stabilized microtubule organization in the oocyte, which is essential for polarization (Fig. 7F).

## Discussion

Cell polarity is a fundamental biological process that drives developmental events across many contexts. Mechanisms of cellular polarization can generally be divided into three primary steps: ***1)*** the symmetry-breaking step, which establishes the initial polarized localization of polarity factor/s from a symmetric or random distribution (e.g.,^61,62^), ***2)*** the polarization step, where a cascade of activities originating from localized polarity factors leads to a polarized state (e.g., through post-translational modifications, transport, etc.,)^62^, and ***3)*** the functional output step on the cell cortex, where the localization of specific components to distinct cortical regions facilitates polarized cellular functions (e.g., localized actin contractility in migration, cytoskeletal anchoring of spindles in stem cell asymmetric division, junction formation in epithelial cells, etc.,).^62^ In oocyte polarity, we previously identified the symmetry-breaking step driven by the bouquet MTOC machinery,^4,19^ which initiates the polarization of the Buc protein.^9^ We also demonstrated that the polarization step is mediated by Buc dependent molecular condensation of Buc RNPs, which is regulated and positioned in the oocyte by microtubules.^10^ At the functional output step on the cell cortex, Macf1 and actin-mediated docking of Buc RNPs to the cortex^15,63^ ensures localization of germplasm and dorsal determinants to the vegetal pole, critical for future embryonic axis formation and germline specification.^16^

Here, we uncover a previously unrecognized feature of oocyte polarity, which operates in parallel to the primary steps of polarization above: the acquisition of polarization competence. This concept refers to the oocyte’s ability to develop the potential to polarize, which includes maintaining an optimal centrosome MTOC and microtubule network that are essential for polarity. Numerous examples describe how polarity factors induce polarized cytoskeletal organizations (reviewed in^61,64^). The reciprocal interactions between cytoskeletal organization and polarity factors, potentially downstream of polarizing signals, have long been postulated^65^. In this work, our findings establish a novel mTOR-Stat3-Stathmin pathway that controls the optimal centrosome MTOC and microtubule organization in the oocyte, regulating polarization. We propose that this pathway confers polarization competence to the oocyte.

While Stat3 is widely recognized for its transcriptional regulation, its direct involvement in cell polarization has not been well established previously. However, evidence suggests Stat3 may also play non-transcriptional roles in other polarity contexts. For instance, in murine embryonic fibroblast migration, Stat3 and Stathmin regulate microtubules independently of transcription, likely contributing to the establishment of polarity and the formation of the cell leading edge.^50,66^ Similarly, in sperm, p-Stat3 localizes to the acrosome—a polarized organelle that forms opposite the flagellum,^67^ suggesting a role for Stat3 in polarized organelle formation and/or function. This raises the intriguing possibility that Stat3’s regulation of microtubules may be widely conserved and that the acquisition of polarization competence could be a common step in cell polarity.

Stat3 and Stathmin have been shown to regulate centrosome clustering, MTOC formation, and microtubule stability in cancer contexts,^50,51,54,68–70^ where they operated at least partially through intricate interactions with master centrosome regulators Aurora A (AurA) and Plk1.^51,69^ Stathmin interactions with AurA were required for centrosome clustering.^69^ Down-regulation of either Stat3 or Plk1 reduced recruitment of γTub for MTOC maturation, but either Stathmin up- or down-regulation decreased Plk1 activity.^51,69^ Our study provides the first evidence for centrosome regulation by Stat3-Stathmin in zebrafish oogenesis. In oogenesis, the centrosome MTOC governs key processes such as oogonial division, bouquet formation at meiosis onset, organization of the cleft microtubule network at pachytene-diplotene stages, and it subsequently disassembles thereafter.^9^ However, the regulation over these dynamic organizations and functions of the centrosome MTOC was unknown. We now demonstrate that Stat3 and Stathmin contribute to centrosome regulation and microtubule stability in zebrafish oocytes. Stat3-Stathmin regulation of the oocyte centrosome MTOC likely involves intricate interactions with key centrosome regulators like Aurora A and Plk1, similar to those shown in cancer, but this requires further studies.

Extending from the centrosome MTOC, the microtubule network in the oocyte is also highly dynamic, changing organization to facilitate polarity. For example, microtubules form perinuclear cables at symmetry breaking^9,19^ and a meshwork in the cleft during polarization^9,10^. The dynamic spatial organization of microtubule cables can be further controlled by Stathmin. Opposite gradients of phosphorylated and non-phosphorylated Stathmin were reported by FRET in cell culture, with denser microtubule organization correlating with phosphorylated Stathmin.^66^ Stathmin down-regulation in another cell line increased looping of microtubules, suggesting the control of the spatial form of cables. It is tempting to speculate that localized Stathmin activity in the oocyte can influence microtubule cable dynamics locally to craft their changing spatial organization during oogenesis.

Our discovery that Stat3 functions as a competence factor for oocyte polarization opens exciting possibilities for understanding cell-cell communication in early oogenesis. Stat3’s established roles in regulating cell-cell communication in multiple developmental processes^42,43,53,59^ suggests it may also be involved in oocyte polarization. Communication between the oocyte and granulosa cells is a hallmark of follicular development in oogenesis,^70–72^ and it is plausible that it is also involved in oocyte polarization. For example, the micropyle (a specialized follicle cell that enables sperm entry in fertilization) forms from differentiating follicle cells specifically at the animal pole of the oocyte,^73,74^ and in *buc* mutants, ectopic micropyles form around the oocyte.^17^ This suggest that vegetally localized factors in the oocyte act (directly or indirectly) to restrict micropyle differentiation in surrounding follicle cells animally, by a yet unknown mechanism. However, whether reciprocally, granulosa cells potentially regulate oocyte polarity is unknown. In this context, identifying Stat3 activating signal/s and their sources offers an interesting area for future investigation.

While we identified and focus on mTOR as an activator of Stat3 in oocyte polarity, mTOR is an intracellular factor that intersects with many cytokines and growth factors^75–77^ that could signal to oocytes. In parallel, additional potential upstream cell-cell communication is suggested by the enrichment of the IL-6 pathway with Stat3 in our transcriptomic data (Fig. S7B). IL-6 is a major activator of Stat3^50,51,54,68,69^; IL-6 activates Jak upstream of Stat3 through the gp130 receptor.^50,51,54,68,69^ Future investigations will determine whether IL-6 is required to control Stat3 in oocyte polarity, whether this occurs via autocrine or paracrine mechanisms, and in the latter case, which cells serve as sender cells.

Our findings that mTOR-Stat3-Stathmin are required for centrosome and microtubule organization in the oocyte demonstrate that this regulation, previously known from cancer and cell culture models, is conserved in a physiological context during animal post-embryonic development. Given the conservation of mTOR, Stat3, and Stathmin across species, it is likely that their mechanistic control over centrosome and microtubule regulation like the one we report, extend to other organisms. For instance, the similarities between the zebrafish and mouse Bb, and its common regulation by microtubules in both species,^9,10,22,25^ suggest that the roles of Stat3 may be conserved in mice. Supporting this hypothesis, p-Stat3 was detected in mice oocytes between E12.5 to PND4,^67^ which overlaps with Bb formation. Further, p-Stat3 localized to microtubules and meiotic spindle poles in mouse oocyte maturation.^78^ p-Stat3 was also detected in human fetal and adult ovaries, in both oocytes and granulosa cells,^79^ indicating yet unknown functions in human oogenesis.

Our findings are consistent with reported non-transcriptional regulation by Stat3, but transcriptional regulation of germline development by Stat3 is also established. In *Drosophila*, Stat3 controls primordial germ cell migration to the gonad, as well as germline- and follicle stem cell maintenance.^41^ In turtles, Stat3 mediates temperature-dependent sex determination, upregulating expression of the oocyte transcription factor *foxl2*, and promoting female fate.^80^ Therefore, Stat3 transcriptional and non-transcriptional regulation are very likely to cooperate in parallel in zebrafish oocytes, and future studies will resolve Stat3 transcriptional regulation in oogenesis. Overall, our findings highlight the importance of Stat3 as a major regulator of the germline across species.

Beyond the roles of mTOR-Stat3-Stathmin, we also present the development of a powerful *ex-vivo* system for studying oogenesis. For the first time in zebrafish, our long-term ovary culture method enables the development of follicles from oogonia in ovaries *ex-vivo*. This process reliably recapitulates key aspects of early oogenesis in the intact ovary, while offering the added advantage of being highly accessible for *ex-vivo* analyses. As demonstrated throughout our work, this system provides an invaluable platform for rapid functional studies, robust candidate screening, and can be combined with genetics to investigate processes that would otherwise be difficult to address. In the future, when combined with live imaging techniques, this system will enable real-time investigation of dynamic cellular processes in specific cells over extended time periods, which holds great promise for advancing oogenesis research.

In conclusion, our study identifies a critical mTOR-Stat3-Stathmin pathway in oocyte polarization, revealing previously unknown regulatory mechanisms essential for microtubule organization and centrosome dynamics. We propose these mechanisms establish the cell’s competence to polarize, uncovering a key regulatory aspect of oocyte polarity. Overall, the uncovered functions of the mTOR-Stat3-Stathmin pathway in oogenesis highlight broader implications in developmental biology, and advance our understanding of female reproduction.

## Resource Availability

### Lead Contact

Requests for further information and resources should be directed to and will be fulfilled by the lead contact, Yaniv M. Elkouby.

### Material Availability

This study did not generate new unique reagents.

### Data and Code Availability

All data are included with this manuscript.

Microscopy data reported in this paper will be shared by the lead contact upon request. Data reported in this paper will be shared by the lead contact upon request.

Any additional information required to reanalyze the data reported in this paper is available from the lead contact upon request.

## Acknowledgements

We are grateful to Lilianna Solnica-Krezel (Washington University in St. Louise School of Medicine) and Sven Reischauer (Max Planck Institute for Heart and Lung Research) for the *stat3^stl^*^27^ line. This work was funded by the Israel Science Foundation – grant no. 558/19 to YME. AS is grateful for the financial support of the Harper Fellowship, from the Neubauer Family Foundation.

## Author contributions

Conceptualization: AS, YB, YME.

Methodology: AS, YB, YME.

Investigation: AS, YB.

Visualization: AS, YB, YME.

Funding acquisition: AS, YME.

Supervision: YME.

Writing – original draft: AS, YB, YME.

## Material and methods

### Ethics statement

All animal experiments were supervised by the Hebrew University Authority for Biological Models, according to the Hebrew University School of Medicine Institutional Animal Care and Use Committee under ethics requests MD-18-15600-2, and MD-20-16228-4, and accredited by the Association for Assessment and Accreditation of Laboratory Animal Care International.

### Fish lines and gonad collections

Juvenile ovaries were collected from 4-6 week post-fertilization (wpf) juvenile fish. Fish had a standard length (SL) measured according to^81^ and were consistently ∼8-12mm. Ovary collection was done as in^9^. Briefly, to fix the ovaries for immunostaining, fish were cut along the ventral midline and the lateral body wall was removed. The head and tail were removed and the trunk pieces, with the exposed abdomen containing the ovaries, were fixed in 4% PFA at 4°C overnight with nutation. Trunks were then washed in PBS and ovaries were finely dissected in cold PBS. Ovaries were washed in PBS and then either stored in PBS at 4°C in the dark, or dehydrated and stored in 100% MeOH at −20°C in the dark. Fish lines used in this research are: *TU* wild type, *stat3^stl^*^27^ ^48,49^.

### Genotyping

Genotyping was performed at 3 wpf, after which fish were rested and raised in the system until the collection of their ovaries at 4-6 wpf. Fish were anaesthetized in 0.02% Tricaine (Sigma Aldrich, #A5040) in system water, and a piece of their fin tail was clipped for DNA extraction. DNA was extracted using the standard *HotSHOT* DNA preparation method. *stat3^stl^*^27^ mutant fish were genotyped by KASP genotyping (LGC, Teddington, UK). KASP-PCR was performed on an Applied Biosystems StepOnePlus machine and following the manufacturer instructions, using the following SNP sequence: ATGCATGTTATAACAATCTGAAAACTGAAAATACTAAAACTGAAACTAATATTTAGCTATTGCTTGGGTATAACCTCTTACTCATCCTCCACAGGACA[TGGAGCA/-]GAAGATGAAAATGTTGGA GAATCTGCAGGATGATTTTGATTTCAACTACAAAACTCTTAAGAGTGCTGGA

### Fluorescence immunohistochemistry (IHC)

Fluorescence immunohistochemistry (IHC) was performed as in^9^. Briefly, ovaries were washed 4 x 20 minutes in PBT (0.3% Triton X-100 in 1xPBS. If stored in MeOH, ovaries were gradually rehydrated before washes. Ovaries were blocked for 1.5-2 hours in blocking solution (10% FBS in PBT) at room temperature, and then incubated with primary antibodies in blocking solution at 4°C overnight. Ovaries were washed 4 x 20 minutes in PBT and incubated with secondary antibodies in fresh blocking solution for 2 hours, and were light protected from this step onward. Ovaries were then washed 4 x 20 minutes in PBT and incubated in PBT containing DAPI (1:1000, Molecular Probes), with or without DiOC6 (1:5000, Molecular Probes) for 50 minutes and washed 2 x 5 minutes in PBT and 2 x 5 minutes in PBS. Ovaries were mounted in Vectashield (with DAPI, Vector Labs) between two #1.5 coverslips using a 120μm spacer (Molecular Probes).

Primary antibodies used were Buc^6^ (1:500; Yenzym Antibodies LLC), αTubulin (1:1000, Merck), γTubulin (1:400, Sigma-Aldrich), εTub (1:200, Sigma-Aldrich), Phospho-Stat3 (Tyr705) (1:200; cell signaling), Phospho-Stathmin (Ser16) (1:200; cell signaling). Secondary antibodies used were Alexa Fluor 488 and 594 (1:500; Molecular Probes). Vital dyes used were: DiOC6 (1:5000, Molecular Probes), Mitotracker (1:2000, Molecular Probes).

### *Ex-vivo* ovary culture

#### Preparation of Ovary Extract

Ovary extracts were prepared as described in Bontems et al. (2009), with modifications. Briefly, ovary extracts were prepared from whole zebrafish ovaries obtained 8 months post-fertilization. The ovaries were disrupted in a medium composed of 60% Leibovitz’s L-15 and 40% Hank’s Balanced Salt Solution (HBSS) by performing three freeze-thaw cycles using liquid nitrogen. Samples were centrifuged at maximum speed (20,000 x g) for 30 minutes at 4°C. The clear supernatant was collected and stored at −80°C at a concentration of one ovary per milliliter in the same 60% L-15/40% HBSS medium.

#### Preparation of the Culture Dishes

Culture dishes were prepared under sterile tissue culture conditions. High glass-bottom 35-mm dishes were coated with poly-D-lysine (PDL) and incubated overnight at room temperature. Dishes were then washed sequentially with 1× PBS, followed by three washes with double-distilled water (ddH₂O). Dishes were further incubated for 2 hours before use. Microseal spacers were cut to size and placed on the prepared plates.

#### Preparation of the Culture Medium

Basic ovary culture medium (BOCM) contained ovary extract (see Preparation of Ovary Extract), fetal bovine serum (FBS, 10%; Sigma), HEPES buffer solution (20 μM; Gibco), L-alanyl-L-glutamine (2 mM; Sigma), penicillin-streptomycin-amphotericin B solution (1%; Sigma), fish serum (1%; Oncorhynchus mykiss serum, purchased from Kibbutz Dan), and insulin-transferrin-selenium (ITS, 1%; Sigma), in Dulbecco’s modified Eagle’s medium (DMEM) high-glucose (Sartorius). Complete ovary culture medium 1 (COCM1) is identical to BOCM, except it also contains 17α,20β-dihydroxy-4-pregnen-3-one (DHP; Cayman Chemical, #16146) at final concentrations of 1 ng ng/mL, and estradiol-17β (E2; Sigma-Aldrich, #E2758) at final concentrations of 1 ng ng/mL. Complete ovary culture medium 2 (COCM2) is identical to BOCM, except it also contains DHP at final concentrations of 1 ng ng/mL.

Ovaries were cultured under a 5% CO₂ atmosphere at 28 °C, and culture medium was refreshed every 2 days over the course of culture. At the end of experiments cultured ovaries were fixed for analyses as above.

### EdU pulse-chase and Click Chemistry

Ovaries were cultured and pulse-labeled with EdU overnight, washed by replacing culture medium, cultured for desired period of times per experiment, and were fixed post-culture. Detection of EdU incorporation into DNA was performed using the iClick™ EdU Andy Fluor™ 594 Imaging Kit (ABP Biosciences, #A005). Ovaries were washed in 3% bovine serum albumin (BSA) in PBS (pH 7.4) for 10 minutes, then washed 4 × 20 minutes in PBT (0.5% Triton® X-100 in PBS). An additional 10-minute wash in 3% BSA was performed before adding the iClick reaction cocktail. The reaction cocktail consisted of 430 μL of iClick EdU Reaction Buffer, 20 μL of CuSO₄, 1.5 μL of Andy Fluor 594 azide, and 50 μL of 1× Reaction Buffer Additive, for a total volume of 500 μL per sample. The cocktail was added to each plate, and ovaries were incubated for 1 hour. After incubation, ovaries were washed sequentially: 1 × 10 minutes in BSA, 2 × 5 minutes in PBT, and 2 × 5 minutes in PBS.

### Drug Treatments in *Ex-Vivo* Zebrafish Ovary Culture

Ovaries in culture were treated with COCM1+2 containing 100 μM Pyrimethamine, or 50 nM Rapamycin, or 75 μM GDP366 or their combinations as described in the text, for 7 days. Ovaries were cultured and collected post-culture as described above.

### Drug Treatments in Whole Live Zebrafish

Juvenile zebrafish at 4 wpf were used for drug treatments in whole living fish. The fish, measuring between 5 and 11 mm in standard length (SL) were kept in fish system water containing one of the following treatments: DMSO, or 100 μM Pyrimethamine (Sigma-Aldrich #46706). The treatments were administered in alternating 2-days cycles of treatment and rest (fish water without DMSO or Pyrimethamine), over the course of two weeks. Fish were normally fed as per our routine husbandry conditions and their water were manually replaced after each feeding with fish water containing either DMSO or Pyrimethamine in a treatment cycle or plane fish water in a rest cycle. At the end of experiments fish SL was measured, and ovaries were collected as described above.

### Confocal microscopy

Images were acquired on a Zeiss LSM 880 confocal microscope using a 40X lense. The acquisition setting was set across samples and experiments to: XY resolution=1104×1104 pixels, 12-bit, 2x sampling averaging, pixel dwell time=0.59sec, zoom=0.8X, pinhole adjusted to 1.1μm of Z thickness, increments between images in stacks were 0.53μm, laser power and gain were set in an antibody-dependent manner to 4-9% and 400-650, respectively, and below saturation condition. Unless otherwise noted, shown images are partial Sum Z-projection. Acquired images were not manipulated in a non-linear manner, and only contrast/brightness and region of interest (ROI) were adjusted in Fiji. All Figures were made using Adobe Photoshop CC 2014 and BioRender.com, in compliance with BioRender’s terms of use.

### Three-dimensional segmentation

Confocal raw data images of entire z-stacks of oocytes were extracted from whole ovary images and imported to IMARIS. Nuclei, centrosomes and Buc condensates were segmented in 3D using blend volume rendering mode, and signal brightness and object transparency were slightly and linearly adjusted in all channels to optimize visualization. Animation frames were made using the key frame animation tool.

### Cell culture

The human epithelial ovarian cancer cell line A2780 was cultured in Dulbecco’s modified Eagle’s medium (DMEM) supplemented with 10% fetal bovine serum (FBS) and antibiotics (100 U/ml penicillin and 100 μg/ml streptomycin). Cells were maintained at 37°C in a humidified atmosphere with 5% CO₂. A2780 cells were treated with pyrimethamine (125 μg/ml) for 24 h, harvested, washed with PBS, and fixed in 4% PFA at 4°C for 24 h. After centrifugation and two additional PBS washes, cells were processed for immunofluorescence. Cells were stained with primary antibody against Cleaved Caspase3(c-Caspase3) (1:400, Cell Signaling Tech). DAPI was used to stain DNA for 30min. Imaging was performed using confocal fluorescence microscope.

### Bioinformatic analysis of oocyte transcriptomic data

Analysis of the RNAseq data was performed as in.^6^ The canonical pathway analysis for genes enriched in different clusters was performed using the Ingenuity Pathway Analysis (IPA) suite (QIAGEN Inc.). Heatmaps of the expression levels of the genes associated with the different pathways were created using Clustvis.^83^

### Statistical Analysis

All statistical analysis and data plotting were performed using the GraphPad Prism 7 software. Datasets were tested with two-way ANOVA. In the figures, P values are indicated by asterisks as follows: *P < 0.05, **P < 0.01, ***P < 0.001, and ****P < 0.0001. ns, not significant at P > 0.05).

## Supplemental information

Figures S1-S7.

Videos S1-S4.

## Supplemental Figure Legends

**Figure S1. Methodology and set up of the ovary culture system. A.** Top: preparation of glass bottom dishes – PDL coating and set up of adherent sealing around the glass. Middle: Ovary dissection – ovaries are dissected from juvenile fish (SL is shown, ruler grid are 1 mm) by cutting along the indicated lines (cyan; Methods). Dissected ovaries (cyan arrowheads) are cleaned from connective tissues and placed on the glass bottom dish. Bottom: Dishes are transferred to sterile conditions and ovaries are covered by a 50 μm cell strainer sieve, which is glued to the adherent sealing around the glass, and medium is added. Right panels show the final set up with a cultured ovary placed on the glass bottom of the dish and covered by the cell strainer sieve in culture media. For A-C, see Methods. **B.** Cultured ovaries in COCM1 are vital through 10 dpc as shown by live-labeling with Mitortracker at 10 dpc. n=4 ovaries. Scale bar is 40 μm.

**Figure S2. Efficient inhibition by Pyrimethamine. A.** Representative images of A2780 cells treated with DMSO or Pyrimethamine and labeled for cCaspas3 (red) and DAPI (blue) show detection of cCaspas3 in Pyrimethamine- but not DMSO-treated cells. Scale bars are 50 μm. **B.** Rates of apoptosis as indicated by Caspas3+ cells in the experiments in A. n=number of cells. Bars are mean±SD.

**Figure S3. Supporting information for Figure 2D-F. A.** SL of fish from which ovaries were collected for culture in the experiments in Fig. 2D-F. Fish of typical and consistent range of SL were used. n=number of fish. **B.** Inhibition of Stat3 *ex-vivo* does not affect viability. Representative images of ovaries post culture treated with either DMSO or Stat3 inhibitor, and live-labeled for Mitotracker show similar normal signals and viability. n= 3 DMSO treated and 3 Stat3 inhibitor treated ovaries. Scale bars are 40 μm. **C.** Inhibition of Stat3 *ex-vivo* does not affect gross ovarian development. Representative overview images of ovaries post culture treated with either DMSO or Stat3 inhibitor, exhibit oocytes at the typical range of developmental stages, as well as normal general morphology. Example oocytes are outlined at oogonia stages (red), as well as primordial follicles at pachytene (cyan), and diplotene (blue) stages. n=12 DMSO treated ovaries and 12 Stat3 inhibitor treated ovaries. Scale bars are: 50 μm.

**Figure S4. Inhibition of Stat3 in whole fish perturbs oocyte polarity. A.** Experimental set up: wt juvenile fish in standard husbandry conditions were treated with cycles of Stat3 inhibitor or DMSO administration (in fish water) and rest, over the course of two weeks between 4 wpf and 6 wpf. At 6 wpf fish were scarified and ovaries were dissected for analyses. **B.** Treatments of DMSO and Stat3 inhibitor do not affect apparent normal post-embryonic development. DMSO and Stat3 inhibitor treated fish have similar SL before treatment and reach similar and normal SL post treatment. n=number of fish. ns=not significant. **C.** Representative images of ovaries from fish treated with DMSO (left) or Stat3 inhibitor (two right panels), co-labeled with DiOC6 (green) and DAPI (blue). Ovaries from DMSO treated fish showed normal gross ovarian development with a typical range of oocyte stages. Ovaries from fish treated with Stat3 inhibitor, either showed normal gross ovarian development (middle panel), or appeared underdeveloped (right panel). Underdeveloped ovaries were thinner and contained fewer oocytes in primordial follicle stages. The distribution of ovarian phenotypes are plotted in the right. n=number of ovaries. Scale bars are 40 μm. **D.** Inhibition of Stat3 in whole fish perturbs Buc polarity. Representative images of oocytes (dashed outlines) at symmetry breaking, early-, and late cleft, and mature Bb stages in ovaries treated with DMSO (top) or Stat3 inhibitor (bottom) and labeled for Buc (red, white arrowheads) and DAPI (greyscale). The distribution of Buc phenotypes per stage is plotted below each panel. n=number of oocytes, from 5 ovaries from fish treated with either DMSO or Stat3 inhibitor each. Scale bars are 10 μm. **E.** Inhibition of Stat3 in whole fish perturbs the polarized localization of Bb-enriched organelles. Representative images of oocytes (dashed outlines) at symmetry breaking, early-, and late cleft, and mature Bb stages in ovaries treated with DMSO (top) or Stat3 inhibitor (bottom) and labeled for DiOC6 (cyan, white arrowheads). The distribution of DiOC6 phenotypes per stage is plotted below each panel. n=number of oocytes, from 6-8 ovaries from fish treated with either DMSO or Stat3 inhibitor each. Scale bars are 10 μm.

**Figure S5. Supporting information for Figure 3. A.** *stat3^-/-^*larvae exhibit scoliosis at 7 dpf, as expected.^48^ **B.** At 22 dpf, *stat3^-/-^* reach SL=7.5 mm, while wt fish reach SL=∼9 mm on average. However, we did not correct for the scoliosis phenotype (A), which could contribute to this slight decrease. n=number of fish. **C.** *stat3^-/-^* fish are lethal at juvenile stages. As previously reported, the proportions of homozygous *stat3^-/-^* fish decrease over time starting from 15 dpf and they are not detected at 45 dpf.^48^ For our experiments, we collected ovaries from fish at 30 dpf, when few homozygous fish remain viable, as shown from genotyping results per cross for 10 representative crosses. We uniformly pooled wt and *stat3^-/-^* fish from several clutches per experiment. n=number of fish.

**Figure S6. Supporting information for** Figure 3C**. A.** Representative images of oocytes at mature Bb stages (dashed outline) from wt (top) and *stat3^-/-^* (bottom) ovaries, co-labeled for Buc (red, white arrowheads) and DAPI (greyscale). n=4 oocytes from wt ovaries and 3 oocytes from *stat3^-/-^* ovaries. Scale bars are 10 μm. **B.** Representative images of oocytes at mature Bb stages (dashed outline) from wt (top) and *stat3^-/-^* (bottom) ovaries, co-labeled for αTub (green) and DAPI (greyscale). n=3 oocytes from wt ovaries and 3 oocytes from *stat3^-/-^*ovaries. Scale bars are 10 μm.

**Figure S7. Supporting information for Figure 7. A.** Expression levels through oogenesis of the genes included in the mTOR pathways from cluster #1. **B.** Genes associated with the Il-6 pathway are expressed in oogenesis. Top: The Il-6 pathways is enriched in cluster #1. Bottom: Expression levels through oogenesis of the genes included in the Il-6 pathways from cluster #1.

## Supplemental Video Legends

**Video S1. Normal general ovarian development in *stat3* mutants.** View through the Z-stack of images in Fig. 3B. Wt (left) and *stat3* (right) ovaries co-labeled for DiOC6 (cyan), Phalloidin (inverted LUT, black), and DAPI (red) are shown. Scale bars are 50 μm.

**Video S2. Microtubule destabilization in *stat3* ovaries.** View through the Z-stack of Wt (left) and *stat3* (right) ovaries co-labeled for αTub (inverted LUT) and DAPI (blue). Scale bars are 50 μm. Images in Fig. 4A are representative oocytes different stages from this experiment.

**Video S3. Centrosome aberrations in *stat3* ovaries.** Videos of the centrosome phenotypes in the representative oocytes from Fig. 4D. γTub is in maroon and DPAI in grey.

**Video S4. Concomitant centrosome and Buc phenotypes in *stat3* ovaries.** Videos of the centrosome and Buc phenotypes in the representative oocytes from Fig. 4F. γTub is in maroon, Buc in cyan, and DPAI in grey.

## References

1. Charalambous, C., Webster, A., and Schuh, M. (2023). Aneuploidy in mammalian oocytes and the impact of maternal ageing. Nat. Rev. Mol. Cell Biol. 24, 27–44. 10.1038/s41580-022-00517-3.

2. Elkouby, Y.M., and Mullins, M.C. (2017). Methods for the analysis of early oogenesis in Zebrafish. Dev. Biol. 430, 310–324. 10.1016/j.ydbio.2016.12.014.

3. Elkouby, Y.M. (2017). All in one - integrating cell polarity, meiosis, mitosis and mechanical forces in early oocyte differentiation in vertebrates. Int. J. Dev. Biol. 61, 179–193. 10.1387/ijdb.170030ye.

4. Mytlis, A., Levy, K., and Elkouby, Y.M. (2023). The many faces of the bouquet centrosome MTOC in meiosis and germ cell development. Curr. Opin. Cell Biol. 81, 102158. 10.1016/j.ceb.2023.102158.

5. Elkouby, Y.M., and Mullins, M.C. (2017). Coordination of cellular differentiation, polarity, mitosis and meiosis - New findings from early vertebrate oogenesis. Dev. Biol. 430, 275–287. 10.1016/j.ydbio.2017.06.029.

6. Bogoch, Y., Jamieson-Lucy, A., Vejnar, C.E., Levy, K., Giraldez, A.J., Mullins, M.C., and Elkouby, Y.M. (2022). Stage specific transcriptomic analysis and database for zebrafish oogenesis. Front. Cell Dev. Biol. 10, 826892. 10.3389/fcell.2022.826892.

7. Langdon, Y.G., and Mullins, M.C. (2011). Maternal and zygotic control of zebrafish dorsoventral axial patterning. Annu. Rev. Genet. 45, 357–377. 10.1146/annurev-genet-110410-132517.

8. Bontems, F., Stein, A., Marlow, F., Lyautey, J., Gupta, T., Mullins, M.C., and Dosch, R. (2009). Bucky ball organizes germ plasm assembly in zebrafish. Curr. Biol. 19, 414–422. 10.1016/j.cub.2009.01.038.

9. Elkouby, Y.M., Jamieson-Lucy, A., and Mullins, M.C. (2016). Oocyte polarization is coupled to the chromosomal bouquet, a conserved polarized nuclear configuration in meiosis. PLoS Biol. 14, e1002335. 10.1371/journal.pbio.1002335.

10. Kar, S., Deis, R., Ahmad, A., Bogoch, Y., Dominitz, A., Shvaizer, G., Sasson, E., Mytlis, A., Ben-Zvi, A., and Elkouby, Y.M. (2025). The Balbiani body is formed by microtubule-controlled molecular condensation of Buc in early oogenesis. Curr. Biol. 35, 315–332.e7. 10.1016/j.cub.2024.11.056.

11. Krishnakumar, P., Riemer, S., Perera, R., Lingner, T., Goloborodko, A., Khalifa, H., Bontems, F., Kaufholz, F., El-Brolosy, M.A., and Dosch, R. (2018). Functional equivalence of germ plasm organizers. PLoS Genet. 14, e1007696. 10.1371/journal.pgen.1007696.

12. Aguero, T., Kassmer, S., Alberio, R., Johnson, A., and King, M.L. (2017). Mechanisms of vertebrate germ cell determination. Adv. Exp. Med. Biol. 953, 383–440. 10.1007/978-3-319-46095-6_8.

13. Jamieson-Lucy, A., and Mullins, M.C. (2019). The vertebrate Balbiani body, germ plasm, and oocyte polarity. Curr. Top. Dev. Biol. 135, 1–34. 10.1016/bs.ctdb.2019.04.003.

14. Jamieson-Lucy, A.H., Kobayashi, M., James Aykit, Y., Elkouby, Y.M., Escobar-Aguirre, M., Vejnar, C.E., Giraldez, A.J., and Mullins, M.C. (2022). A proteomics approach identifies novel resident zebrafish Balbiani body proteins Cirbpa and Cirbpb. Dev. Biol. 484, 1–11. 10.1016/j.ydbio.2022.01.006.

15. Escobar-Aguirre, M., Zhang, H., Jamieson-Lucy, A., and Mullins, M.C. (2017). Microtubule-actin crosslinking factor 1 (Macf1) domain function in Balbiani body dissociation and nuclear positioning. PLoS Genet. 13, e1006983. 10.1371/journal.pgen.1006983.

16. Escobar-Aguirre, M., Elkouby, Y.M., and Mullins, M.C. (2017). Localization in oogenesis of maternal regulators of embryonic development. Adv. Exp. Med. Biol. 953, 173–207. 10.1007/978-3-319-46095-6_5.

17. Marlow, F.L., and Mullins, M.C. (2008). Bucky ball functions in Balbiani body assembly and animal-vegetal polarity in the oocyte and follicle cell layer in zebrafish. Dev. Biol. 321, 40–50. 10.1016/j.ydbio.2008.05.557.

18. Dosch, R., Wagner, D.S., Mintzer, K.A., Runke, G., Wiemelt, A.P., and Mullins, M.C. (2004). Maternal control of vertebrate development before the midblastula transition: mutants from the zebrafish I. Dev. Cell 6, 771–780. 10.1016/j.devcel.2004.05.002.

19. Mytlis, A., Kumar, V., Qiu, T., Deis, R., Hart, N., Levy, K., Masek, M., Shawahny, A., Ahmad, A., Eitan, H., et al. (2022). Control of meiotic chromosomal bouquet and germ cell morphogenesis by the zygotene cilium. Science 376, eabh3104. 10.1126/science.abh3104.

20. Kloc, M., Bilinski, S., and Etkin, L.D. (2004). The Balbiani body and germ cell determinants: 150 years later. Curr. Top. Dev. Biol. 59, 1–36. 10.1016/S0070-2153(04)59001-4.

21. Amin, R., Bukulmez, O., and Woodruff, J.B. (2023). Visualization of Balbiani Body disassembly during human primordial follicle activation. MicroPubl. Biol. 2023. 10.17912/micropub.biology.000989.

22. Lei, L., and Spradling, A.C. (2016). Mouse oocytes differentiate through organelle enrichment from sister cyst germ cells. Science 352, 95–99. 10.1126/science.aad2156.

23. Pepling, M.E., Wilhelm, J.E., O’Hara, A.L., Gephardt, G.W., and Spradling, A.C. (2007). Mouse oocytes within germ cell cysts and primordial follicles contain a Balbiani body. Proc Natl Acad Sci USA 104, 187–192. 10.1073/pnas.0609923104.

24. Hertig, A.T. (1968). The primary human oocyte: some observations on the fine structure of Balbiani’s vitelline body and the origin of the annulate lamellae. Am. J. Anat. 122, 107–137. 10.1002/aja.1001220107.

25. Niu, W., and Spradling, A.C. (2022). Mouse oocytes develop in cysts with the help of nurse cells. Cell 185, 2576–2590.e12. 10.1016/j.cell.2022.05.001.

26. Tworzydlo, W., Marek, M., Kisiel, E., and Bilinski, S.M. (2017). Meiosis, Balbiani body and early asymmetry of Thermobia oocyte. Protoplasma 254, 649–655. 10.1007/s00709-016-0978-7.

27. Bose, M., Lampe, M., Mahamid, J., and Ephrussi, A. (2022). Liquid-to-solid phase transition of oskar ribonucleoprotein granules is essential for their function in Drosophila embryonic development. Cell 185, 1308–1324.e23. 10.1016/j.cell.2022.02.022.

28. Eichler, C.E., Li, H., Grunberg, M.E., and Gavis, E.R. (2023). Localization of oskar mRNA by agglomeration in ribonucleoprotein granules. PLoS Genet. 19, e1010877. 10.1371/journal.pgen.1010877.

29. Lerit, D.A., and Gavis, E.R. (2011). Transport of germ plasm on astral microtubules directs germ cell development in Drosophila. Curr. Biol. 21, 439–448. 10.1016/j.cub.2011.01.073.

30. Becalska, A.N., and Gavis, E.R. (2010). Bazooka regulates microtubule organization and spatial restriction of germ plasm assembly in the Drosophila oocyte. Dev. Biol. 340, 528–538. 10.1016/j.ydbio.2010.02.006.

31. Fuentes, R., Marlow, F.L., Abrams, E.W., Zhang, H., Kobayashi, M., Gupta, T., Kapp, L.D., DiNardo, Z., Heller, R., Cisternas, R., et al. (2024). Maternal regulation of the vertebrate oocyte-to-embryo transition. PLoS Genet. 20, e1011343. 10.1371/journal.pgen.1011343.

32. Wagner, D.S., Dosch, R., Mintzer, K.A., Wiemelt, A.P., and Mullins, M.C. (2004). Maternal control of development at the midblastula transition and beyond: mutants from the zebrafish II. Dev. Cell 6, 781–790. 10.1016/j.devcel.2004.04.001.

33. Ahmad, A., Bogoch, Y., Shvaizer, G., Guler, N., Levy, K., and Elkouby, Y.M. (2025). The piRNA protein Asz1 is essential for germ cell and gonad development in zebrafish and exhibits differential necessities in distinct types of germ granules. PLoS Genet. 21, e1010868. 10.1371/journal.pgen.1010868.

34. Takemoto, K., Imai, Y., Saito, K., Kawasaki, T., Carlton, P.M., Ishiguro, K.-I., and Sakai, N. (2020). Sycp2 is essential for synaptonemal complex assembly, early meiotic recombination and homologous pairing in zebrafish spermatocytes. PLoS Genet. 16, e1008640. 10.1371/journal.pgen.1008640.

35. Blokhina, Y.P., Frees, M.A., Nguyen, A., Sharifi, M., Chu, D.B., Bispo, K., Olaya, I., Draper, B.W., and Burgess, S.M. (2021). Rad21l1 cohesin subunit is dispensable for spermatogenesis but not oogenesis in zebrafish. PLoS Genet. 17, e1009127. 10.1371/journal.pgen.1009127.

36. Rodríguez-Marí, A., Cañestro, C., Bremiller, R.A., Nguyen-Johnson, A., Asakawa, K., Kawakami, K., and Postlethwait, J.H. (2010). Sex reversal in zebrafish fancl mutants is caused by Tp53-mediated germ cell apoptosis. PLoS Genet. 6, e1001034. 10.1371/journal.pgen.1001034.

37. Kumar, V., and Elkouby, Y.M. (2023). Tools to analyze the organization and formation of the germline cyst in zebrafish oogenesis. Development 150. 10.1242/dev.201349.

38. Mytlis, A., and Elkouby, Y.M. (2021). Live and Time-Lapse Imaging of Early Oogenesis and Meiotic Chromosomal Dynamics in Cultured Juvenile Zebrafish Ovaries. Methods Mol. Biol. 2218, 137–155. 10.1007/978-1-0716-0970-5_12.

39. Miura, C., Higashino, T., and Miura, T. (2007). A progestin and an estrogen regulate early stages of oogenesis in fish. Biol. Reprod. 77, 822–828. 10.1095/biolreprod.107.061408.

40. Chen, Y., Jefferson, W.N., Newbold, R.R., Padilla-Banks, E., and Pepling, M.E. (2007). Estradiol, progesterone, and genistein inhibit oocyte nest breakdown and primordial follicle assembly in the neonatal mouse ovary in vitro and in vivo. Endocrinology 148, 3580–3590. 10.1210/en.2007-0088.

41. Herrera, S.C., and Bach, E.A. (2019). JAK/STAT signaling in stem cells and regeneration: from Drosophila to vertebrates. Development 146. 10.1242/dev.167643.

42. Hillmer, E.J., Zhang, H., Li, H.S., and Watowich, S.S. (2016). STAT3 signaling in immunity. Cytokine Growth Factor Rev. 31, 1–15. 10.1016/j.cytogfr.2016.05.001.

43. Johnson, D.E., O’Keefe, R.A., and Grandis, J.R. (2018). Targeting the IL-6/JAK/STAT3 signalling axis in cancer. Nat. Rev. Clin. Oncol. 15, 234–248. 10.1038/nrclinonc.2018.8.

44. Yu, H., Lee, H., Herrmann, A., Buettner, R., and Jove, R. (2014). Revisiting STAT3 signalling in cancer: new and unexpected biological functions. Nat. Rev. Cancer 14, 736– 746. 10.1038/nrc3818.

45. Heppler, L.N., Attarha, S., Persaud, R., Brown, J.I., Wang, P., Petrova, B., Tošić, I., Burton, F.B., Flamand, Y., Walker, S.R., et al. (2022). The antimicrobial drug pyrimethamine inhibits STAT3 transcriptional activity by targeting the enzyme dihydrofolate reductase. J. Biol. Chem. 298, 101531. 10.1016/j.jbc.2021.101531.

46. Liu, Y., Zhou, H., Yi, T., and Wang, H. (2019). Pyrimethamine exerts significant antitumor effects on human ovarian cancer cells both in vitro and in vivo. Anticancer Drugs 30, 571–578. 10.1097/CAD.0000000000000740.

47. Maack, G., and Segner, H. (2003). Morphological development of the gonads in zebrafish. J. Fish Biol. 62, 895–906. 10.1046/j.1095-8649.2003.00074.x.

48. Liu, Y., Sepich, D.S., and Solnica-Krezel, L. (2017). Stat3/Cdc25a-dependent cell proliferation promotes embryonic axis extension during zebrafish gastrulation. PLoS Genet. 13, e1006564. 10.1371/journal.pgen.1006564.

49. Allanki, S., Strilic, B., Scheinberger, L., Onderwater, Y.L., Marks, A., Günther, S., Preussner, J., Kikhi, K., Looso, M., Stainier, D.Y.R., et al. (2021). Interleukin-11 signaling promotes cellular reprogramming and limits fibrotic scarring during tissue regeneration. Sci. Adv. 7, eabg6497. 10.1126/sciadv.abg6497.

50. Ng, D.C.H., Lin, B.H., Lim, C.P., Huang, G., Zhang, T., Poli, V., and Cao, X. (2006). Stat3 regulates microtubules by antagonizing the depolymerization activity of stathmin. J. Cell Biol. 172, 245–257. 10.1083/jcb.200503021.

51. Morris, E.J., Kawamura, E., Gillespie, J.A., Balgi, A., Kannan, N., Muller, W.J., Roberge, M., and Dedhar, S. (2017). Stat3 regulates centrosome clustering in cancer cells via Stathmin/PLK1. Nat. Commun. 8, 15289. 10.1038/ncomms15289.

52. Shen, S., Niso-Santano, M., Adjemian, S., Takehara, T., Malik, S.A., Minoux, H., Souquere, S., Mariño, G., Lachkar, S., Senovilla, L., et al. (2012). Cytoplasmic STAT3 represses autophagy by inhibiting PKR activity. Mol. Cell 48, 667–680. 10.1016/j.molcel.2012.09.013.

53. Wegrzyn, J., Potla, R., Chwae, Y.-J., Sepuri, N.B.V., Zhang, Q., Koeck, T., Derecka, M., Szczepanek, K., Szelag, M., Gornicka, A., et al. (2009). Function of mitochondrial Stat3 in cellular respiration. Science 323, 793–797. 10.1126/science.1164551.

54. Chauvin, S., and Sobel, A. (2015). Neuronal stathmins: a family of phosphoproteins cooperating for neuronal development, plasticity and regeneration. Prog. Neurobiol. 126, 1–18. 10.1016/j.pneurobio.2014.09.002.

55. Fu, J., Hagan, I.M., and Glover, D.M. (2015). The centrosome and its duplication cycle. Cold Spring Harb. Perspect. Biol. 7, a015800. 10.1101/cshperspect.a015800.

56. Gönczy, P. (2015). Centrosomes and cancer: revisiting a long-standing relationship. Nat. Rev. Cancer 15, 639–652. 10.1038/nrc3995.

57. Stathatos, G.G., Dunleavy, J.E.M., Zenker, J., and O’Bryan, M.K. (2021). Delta and epsilon tubulin in mammalian development. Trends Cell Biol. 31, 774–787. 10.1016/j.tcb.2021.03.010.

58. Shi, X., Wang, D., Ding, K., Lu, Z., Jin, Y., Zhang, J., and Pan, J. (2010). GDP366, a novel small molecule dual inhibitor of survivin and Op18, induces cell growth inhibition, cellular senescence and mitotic catastrophe in human cancer cells. Cancer Biol. Ther. 9, 640– 650. 10.4161/cbt.9.8.11269.

59. Akira, S. (2000). Roles of STAT3 defined by tissue-specific gene targeting. Oncogene 19, 2607–2611. 10.1038/sj.onc.1203478.

60. Wilson, M.L., Romano, S.N., Khatri, N., Aharon, D., Liu, Y., Kaufman, O.H., Draper, B.W., and Marlow, F.L. (2024). Rbpms2 promotes female fate upstream of the nutrient sensing Gator2 complex component Mios. Nat. Commun. 15, 5248. 10.1038/s41467-024-49613-2.

61. Campanale, J.P., Sun, T.Y., and Montell, D.J. (2017). Development and dynamics of cell polarity at a glance. J. Cell Sci. 130, 1201–1207. 10.1242/jcs.188599.

62. Ierushalmi, N., and Keren, K. (2021). Cytoskeletal symmetry breaking in animal cells. Curr. Opin. Cell Biol. 72, 91–99. 10.1016/j.ceb.2021.07.003.

63. Gupta, T., Marlow, F.L., Ferriola, D., Mackiewicz, K., Dapprich, J., Monos, D., and Mullins, M.C. (2010). Microtubule actin crosslinking factor 1 regulates the Balbiani body and animal-vegetal polarity of the zebrafish oocyte. PLoS Genet. 6, e1001073. 10.1371/journal.pgen.1001073.

64. Raman, R., Pinto, C.S., and Sonawane, M. (2018). Polarized organization of the cytoskeleton: regulation by cell polarity proteins. J. Mol. Biol. 430, 3565–3584. 10.1016/j.jmb.2018.06.028.

65. Glotzer, M., and Hyman, A.A. (1995). Cell polarity. The importance of being polar. Curr. Biol. 5, 1102–1105. 10.1016/s0960-9822(95)00221-1.

66. Niethammer, P., Bastiaens, P., and Karsenti, E. (2004). Stathmin-tubulin interaction gradients in motile and mitotic cells. Science 303, 1862–1866. 10.1126/science.1094108.

67. Murphy, K., Carvajal, L., Medico, L., and Pepling, M. (2005). Expression of Stat3 in germ cells of developing and adult mouse ovaries and testes. Gene Expr. Patterns 5, 475–482. 10.1016/j.modgep.2004.12.007.

68. Metge, B., Ofori-Acquah, S., Stevens, T., and Balczon, R. (2004). Stat3 activity is required for centrosome duplication in chinese hamster ovary cells. J. Biol. Chem. 279, 41801– 41806. 10.1074/jbc.M407094200.

69. Silva, V.C., and Cassimeris, L. (2013). Stathmin and microtubules regulate mitotic entry in HeLa cells by controlling activation of both Aurora kinase A and Plk1. Mol. Biol. Cell 24, 3819–3831. 10.1091/mbc.E13-02-0108.

70. Webb, R., Buratini, J., Hernandez-Medrano, J.H., Gutierrez, C.G., and Campbell, B.K. (2016). Follicle development and selection: past, present and future. Anim. Reprod. 13, 234–249. 10.21451/1984-3143-AR883.

71. Zhang, H., and Liu, K. (2015). Cellular and molecular regulation of the activation of mammalian primordial follicles: somatic cells initiate follicle activation in adulthood. Hum. Reprod. Update 21, 779–786. 10.1093/humupd/dmv037.

72. Coticchio, G., Dal Canto, M., Mignini Renzini, M., Guglielmo, M.C., Brambillasca, F., Turchi, D., Novara, P.V., and Fadini, R. (2015). Oocyte maturation: gamete-somatic cells interactions, meiotic resumption, cytoskeletal dynamics and cytoplasmic reorganization. Hum. Reprod. Update 21, 427–454. 10.1093/humupd/dmv011.

73. Dingare, C., Niedzwetzki, A., Klemmt, P.A., Godbersen, S., Fuentes, R., Mullins, M.C., and Lecaudey, V. (2018). The Hippo pathway effector Taz is required for cell morphogenesis and fertilization in zebrafish. Development 145. 10.1242/dev.167023.

74. Yi, X., Yu, J., Ma, C., Dong, G., Shi, W., Li, H., Li, L., Luo, L., Sampath, K., Ruan, H., et al. (2019). The effector of Hippo signaling, Taz, is required for formation of the micropyle and fertilization in zebrafish. PLoS Genet. 15, e1007408. 10.1371/journal.pgen.1007408.

75. Meng, D., Frank, A.R., and Jewell, J.L. (2018). mTOR signaling in stem and progenitor cells. Development 145. 10.1242/dev.152595.

76. Ben-Sahra, I., and Manning, B.D. (2017). mTORC1 signaling and the metabolic control of cell growth. Curr. Opin. Cell Biol. 45, 72–82. 10.1016/j.ceb.2017.02.012.

77. Albert, V., and Hall, M.N. (2015). mTOR signaling in cellular and organismal energetics. Curr. Opin. Cell Biol. 33, 55–66. 10.1016/j.ceb.2014.12.001.

78. Haraguchi, S., Ikeda, M., Akagi, S., and Hirao, Y. (2020). Dynamic Changes in pStat3 are Involved in Meiotic Spindle Assembly in Mouse Oocytes. Int. J. Mol. Sci. 21. 10.3390/ijms21041220.

79. Frost, E.R., Ford, E.A., Peters, A.E., Reed, N.L., McLaughlin, E.A., Baker, M.A., Lovell-Badge, R., and Sutherland, J.M. (2020). Signal transducer and activator of transcription (STAT) 1 and STAT3 are expressed in the human ovary and have Janus kinase 1- independent functions in the COV434 human granulosa cell line. Reprod. Fertil. Dev. 32, 1027–1039. 10.1071/RD20098.

80. Wu, P., Wang, X., Ge, C., Jin, L., Ding, Z., Liu, F., Zhang, J., Gao, F., and Du, W. (2024). pSTAT3 activation of Foxl2 initiates the female pathway underlying temperature-dependent sex determination. Proc Natl Acad Sci USA 121, e2401752121. 10.1073/pnas.2401752121.

81. Parichy, D.M., Elizondo, M.R., Mills, M.G., Gordon, T.N., and Engeszer, R.E. (2009). Normal table of postembryonic zebrafish development: staging by externally visible anatomy of the living fish. Dev. Dyn. 238, 2975–3015. 10.1002/dvdy.22113.

82. Riemer, S., Bontems, F., Krishnakumar, P., Gömann, J., and Dosch, R. (2015). A functional Bucky ball-GFP transgene visualizes germ plasm in living zebrafish. Gene Expr. Patterns 18, 44–52. 10.1016/j.gep.2015.05.003.

83. Metsalu, T., and Vilo, J. (2015). ClustVis: a web tool for visualizing clustering of multivariate data using Principal Component Analysis and heatmap. Nucleic Acids Res. 43, W566–70. 10.1093/nar/gkv468.

