## Supplemental Figures and Legends for "A Novel mTOR-Stat3-Stathmin Pathway Establishes Oocyte Polarization Competence by Orchestrating Centrosome Regulation and Microtubule Organization"

**Document S1 – Supplementary Figures and legends**

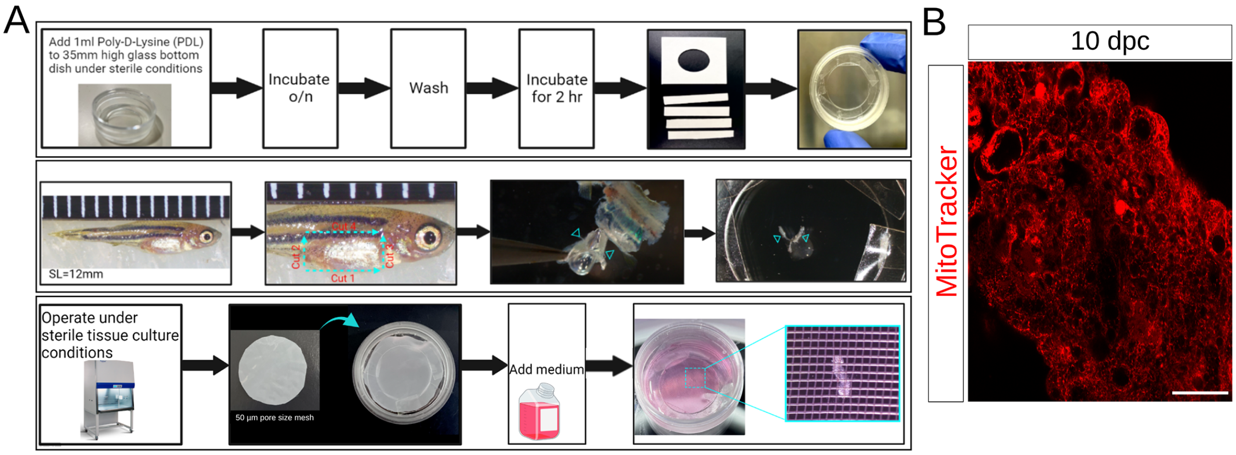


**Figure S1. Methodology and set up of the ovary culture system. A.** Top: preparation of glass bottom dishes – PDL coating and set up of adherent sealing around the glass. Middle: Ovary dissection – ovaries are dissected from juvenile fish (SL is shown, ruler grid are 1 mm) by cutting along the indicated lines (cyan; Methods). Dissected ovaries (cyan arrowheads) are cleaned from connective tissues and placed on the glass bottom dish. Bottom: Dishes are transferred to sterile conditions and ovaries are covered by a 50 μm cell strainer sieve, which is glued to the adherent sealing around the glass, and medium is added. Right panels show the final set up with a cultured ovary placed on the glass bottom of the dish and covered by the cell strainer sieve in culture media. For A-C, see Methods. **B.** Cultured ovaries in COCM1 are vital through 10 dpc as shown by live-labeling with Mitortracker at 10 dpc. n=4 ovaries. Scale bar is 40 μm.

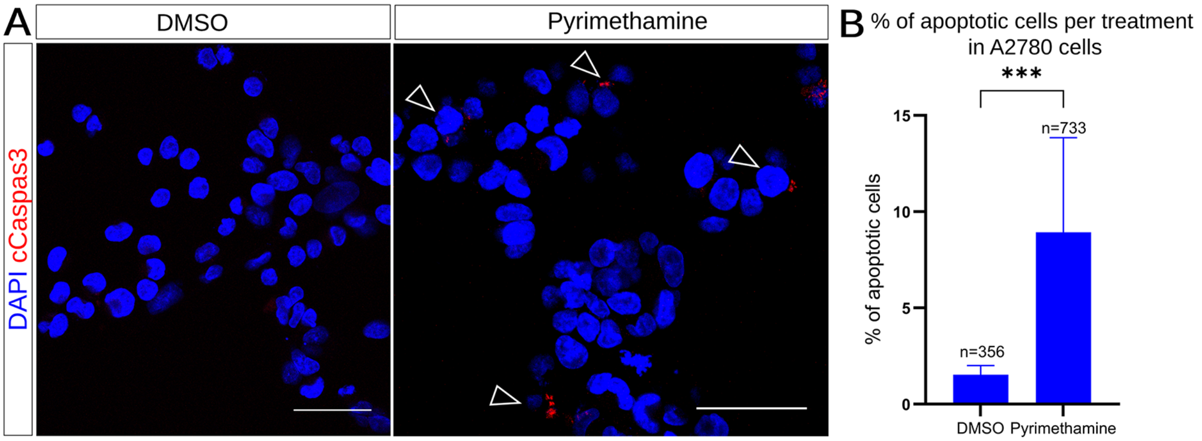


**Figure S2. Efficient inhibition by Pyrimethamine. A.** Representative images of A2780 cells treated with DMSO or Pyrimethamine and labeled for cCaspas3 (red) and DAPI (blue) show detection of cCaspas3 in Pyrimethamine- but not DMSO-treated cells. Scale bars are 50 μm. **B.** Rates of apoptosis as indicated by Caspas3+ cells in the experiments in A. n=number of cells. Bars are mean±SD.

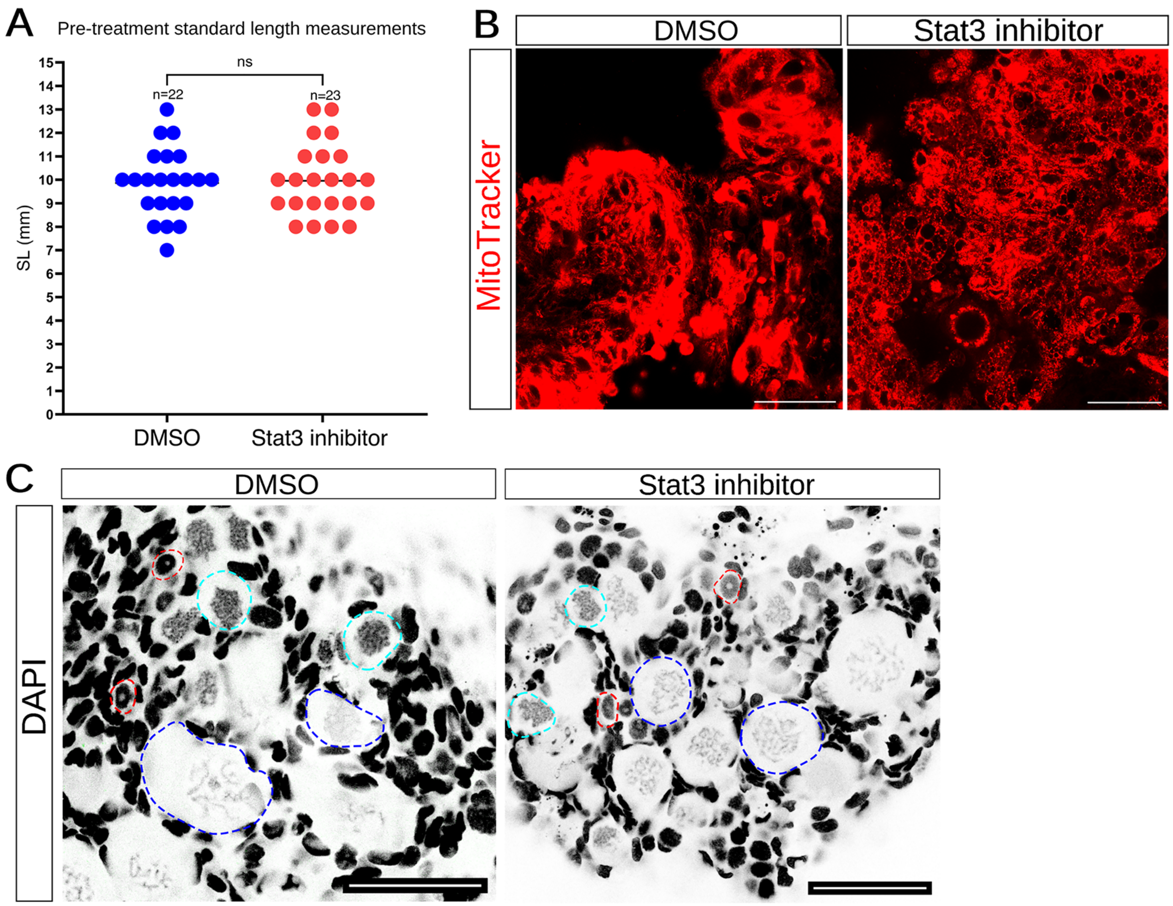


**Figure S3. Supporting information for Figure 2D-F. A.** SL of fish from which ovaries were collected for culture in the experiments in Fig. 2D-F. Fish of typical and consistent range of SL were used. n=number of fish. **B.** Inhibition of Stat3 *ex-vivo* does not affect viability. Representative images of ovaries post culture treated with either DMSO or Stat3 inhibitor, and live-labeled for Mitotracker show similar normal signals and viability. n= 3 DMSO treated and 3 Stat3 inhibitor treated ovaries. Scale bars are 40 μm. **C.** Inhibition of Stat3 *ex-vivo* does not affect gross ovarian development. Representative overview images of ovaries post culture treated with either DMSO or Stat3 inhibitor, exhibit oocytes at the typical range of developmental stages, as well as normal general morphology. Example oocytes are outlined at oogonia stages (red), as well as primordial follicles at pachytene (cyan), and diplotene (blue) stages. n=12 DMSO treated ovaries and 12 Stat3 inhibitor treated ovaries. Scale bars are: 50 μm.

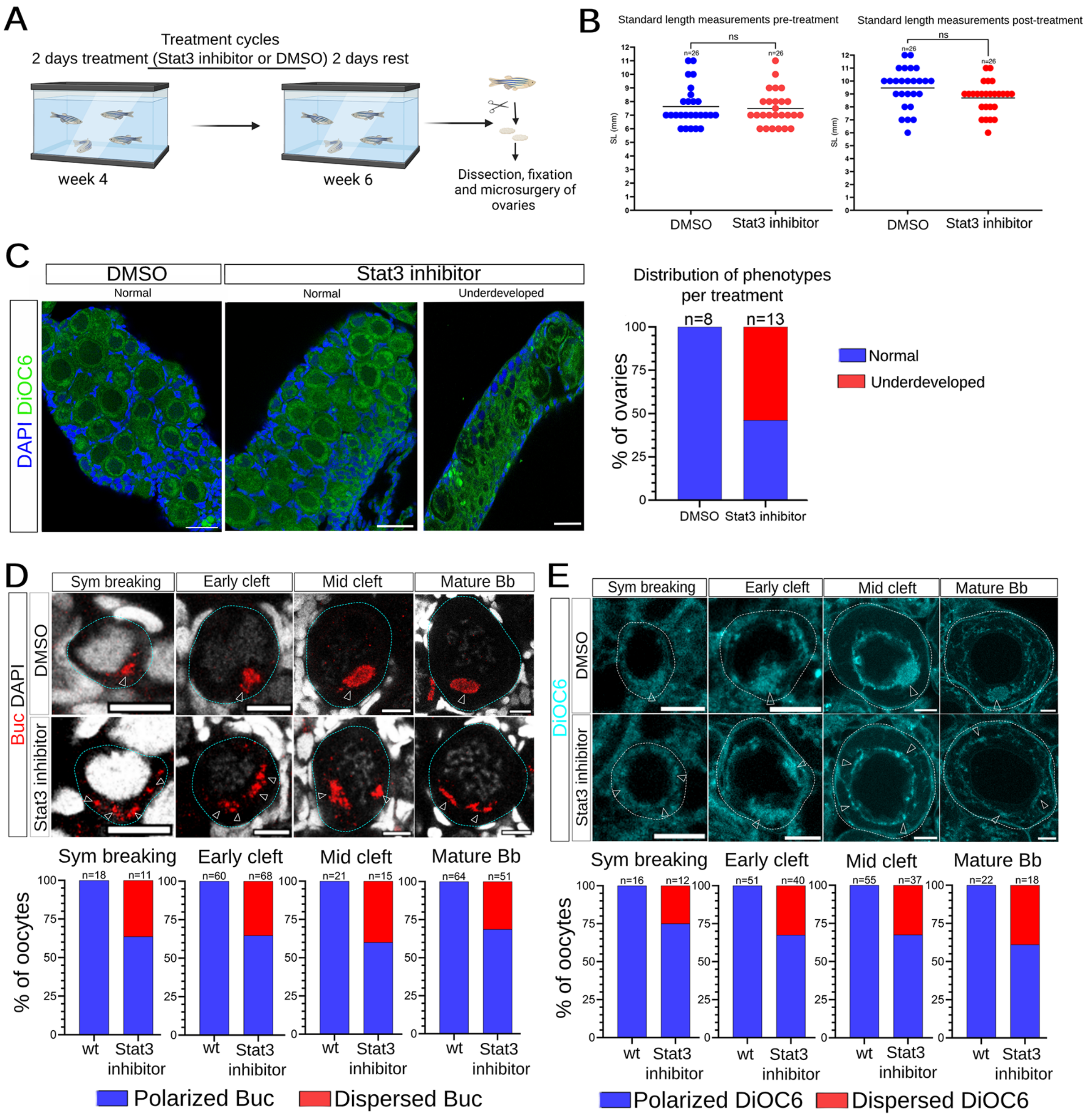


**Figure S4. Inhibition of Stat3 in whole fish perturbs oocyte polarity. A.** Experimental set up: wt juvenile fish in standard husbandry conditions were treated with cycles of Stat3 inhibitor or DMSO administration (in fish water) and rest, over the course of two weeks between 4 wpf and 6 wpf. At 6 wpf fish were scarified and ovaries were dissected for analyses. **B.** Treatments of DMSO and Stat3 inhibitor do not affect apparent normal post-embryonic development. DMSO and Stat3 inhibitor treated fish have similar SL before treatment and reach similar and normal SL post treatment. n=number of fish. ns=not significant. **C.** Representative images of ovaries from fish treated with DMSO (left) or Stat3 inhibitor (two right panels), co-labeled with DiOC6 (green) and DAPI (blue). Ovaries from DMSO treated fish showed normal gross ovarian development with a typical range of oocyte stages. Ovaries from fish treated with Stat3 inhibitor, either showed normal gross ovarian development (middle panel), or appeared underdeveloped (right panel). Underdeveloped ovaries were thinner and contained fewer oocytes in primordial follicle stages. The distribution of ovarian phenotypes are plotted in the right. n=number of ovaries. Scale bars are 40 μm. **D.** Inhibition of Stat3 in whole fish perturbs Buc polarity. Representative images of oocytes (dashed outlines) at symmetry breaking, early-, and late cleft, and mature Bb stages in ovaries treated with DMSO (top) or Stat3 inhibitor (bottom) and labeled for Buc (red, white arrowheads) and DAPI (greyscale). The distribution of Buc phenotypes per stage is plotted below each panel. n=number of oocytes, from 5 ovaries from fish treated with either DMSO or Stat3 inhibitor each. Scale bars are 10 μm. **E.** Inhibition of Stat3 in whole fish perturbs the polarized localization of Bb-enriched organelles. Representative images of oocytes (dashed outlines) at symmetry breaking, early-, and late cleft, and mature Bb stages in ovaries treated with DMSO (top) or Stat3 inhibitor (bottom) and labeled for DiOC6 (cyan, white arrowheads). The distribution of DiOC6 phenotypes per stage is plotted below each panel. n=number of oocytes, from 6-8 ovaries from fish treated with either DMSO or Stat3 inhibitor each. Scale bars are 10 μm.

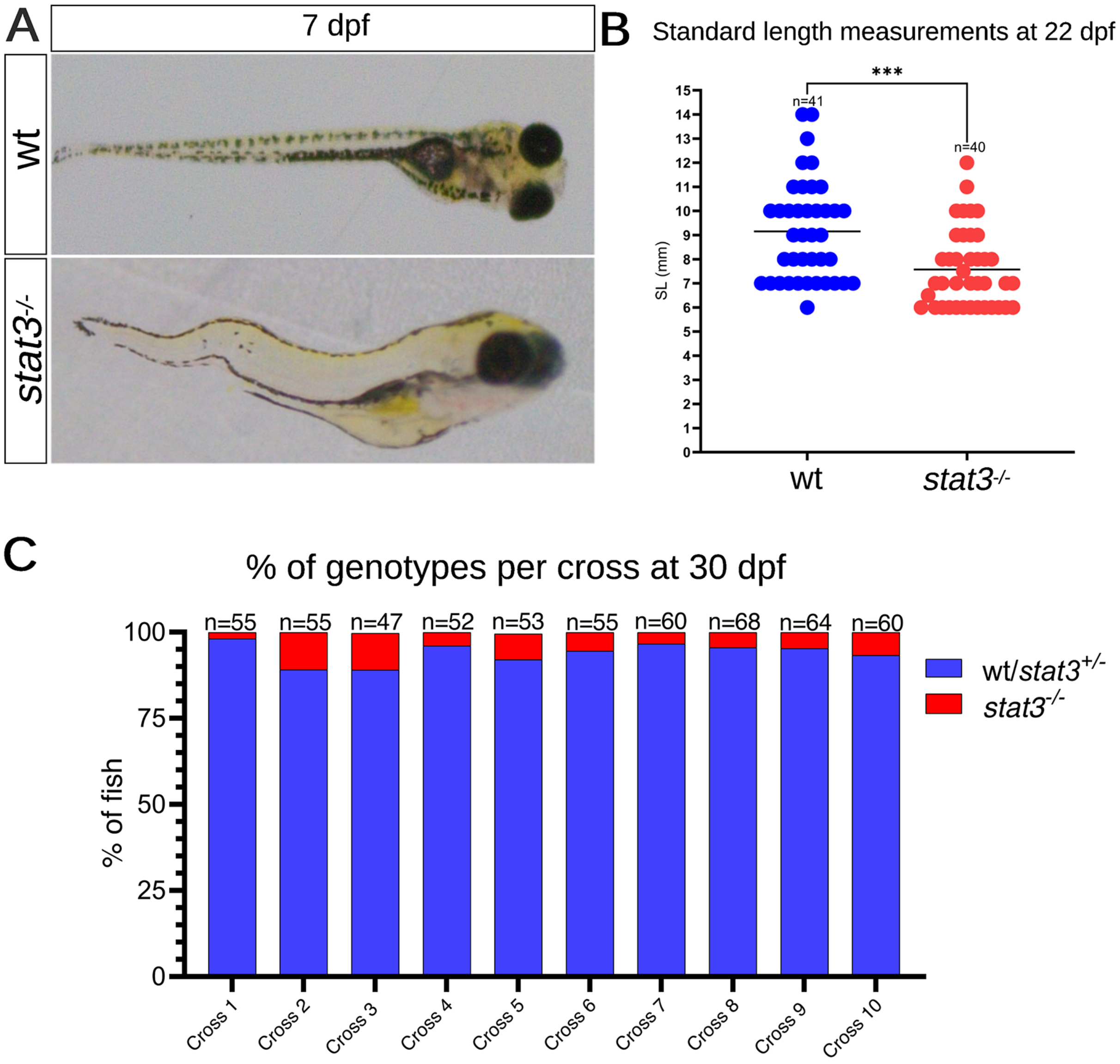


**Figure S5. Supporting information for Figure 3. A.** *stat3*$^{-/-}$ larvae exhibit scoliosis at 7 dpf, as expected.[48] **B.** At 22 dpf, *stat3*$^{-/-}$ reach SL=7.5 mm, while wt fish reach SL=~9 mm on average. However, we did not correct for the scoliosis phenotype (A), which could contribute to this slight decrease. n=number of fish. **C.** *stat3*$^{-/-}$ fish are lethal at juvenile stages. As previously reported, the proportions of homozygous *stat3*$^{-/-}$ fish decrease over time starting from 15 dpf and they are not detected at 45 dpf.[48] For our experiments, we collected ovaries from fish at 30 dpf, when few homozygous fish remain viable, as shown from genotyping results per cross for 10 representative crosses. We uniformly pooled wt and *stat3*$^{-/-}$ fish from several clutches per experiment. n=number of fish.

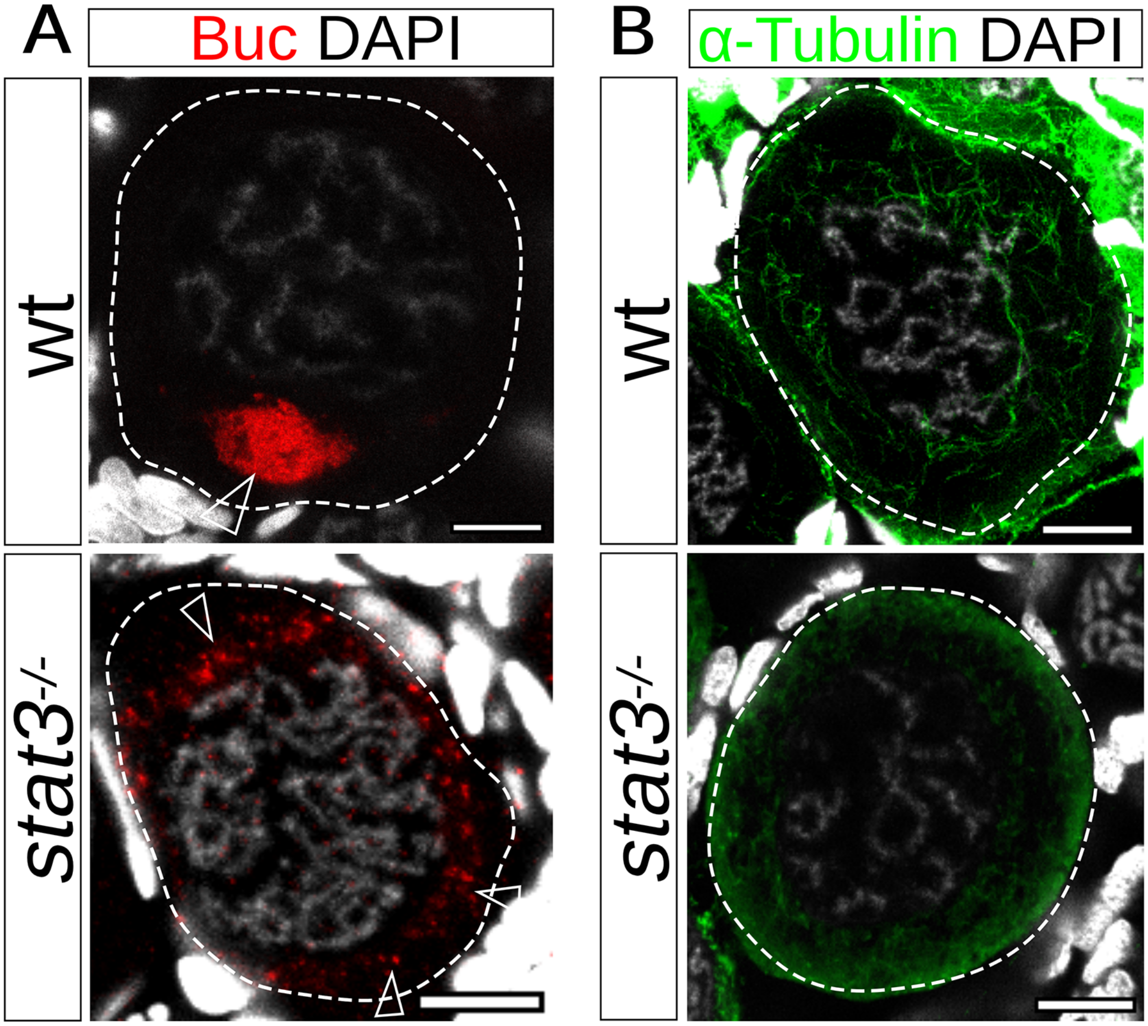


**Figure S6. Supporting information for Figure 3C. A.** Representative images of oocytes at mature Bb stages (dashed outline) from wt (top) and *stat3*$^{-/-}$ (bottom) ovaries, co-labeled for Buc (red, white arrowheads) and DAPI (greyscale). n=4 oocytes from wt ovaries and 3 oocytes from *stat3*$^{-/-}$ ovaries. Scale bars are 10 μm. **B.** Representative images of oocytes at mature Bb stages (dashed outline) from wt (top) and *stat3*$^{-/-}$ (bottom) ovaries, co-labeled for αTub (green) and DAPI (greyscale). n=3 oocytes from wt ovaries and 3 oocytes from *stat3*$^{-/-}$ ovaries. Scale bars are 10 μm.

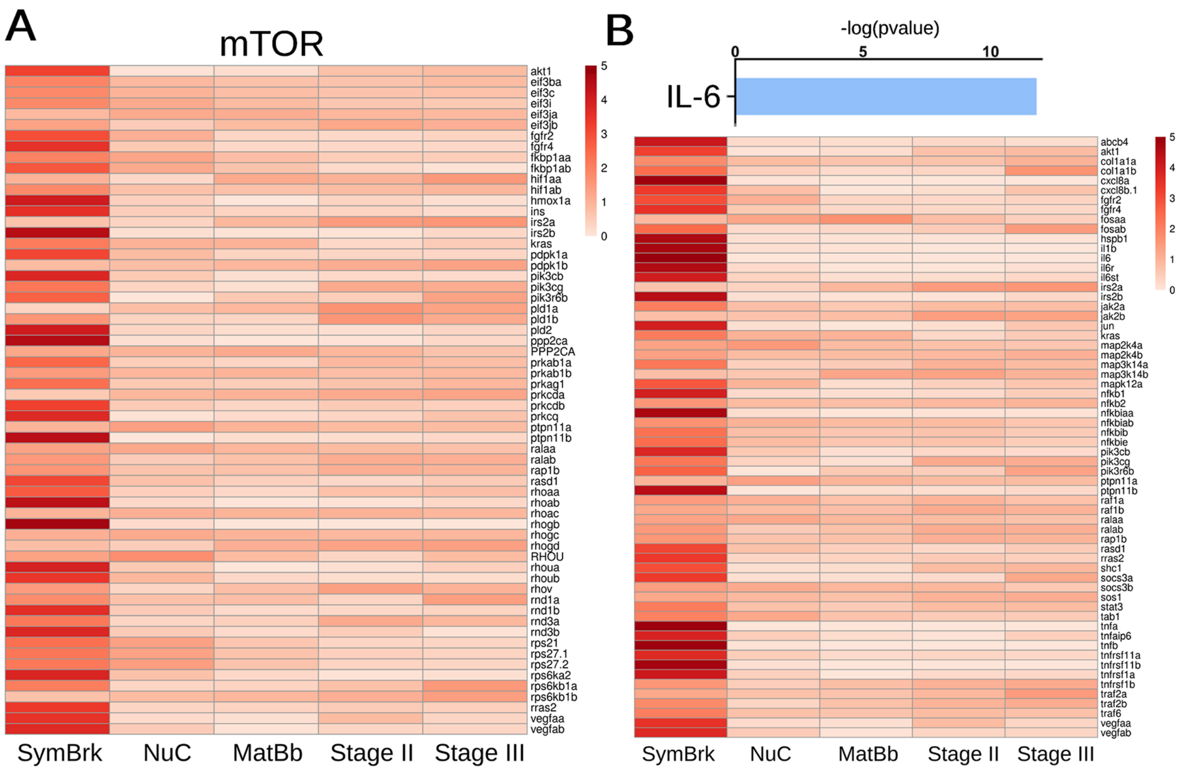


**Figure S7. Supporting information for Figure 7. A.** Expression levels through oogenesis of the genes included in the mTOR pathways from cluster #1. **B.** Genes associated with the Il-6 pathway are expressed in oogenesis. Top: The Il-6 pathways is enriched in cluster #1. Bottom: Expression levels through oogenesis of the genes included in the Il-6 pathways from cluster #1.
